# NBS1-CtIP–Mediated DNA End Resection Regulates cGAS Binding to Micronuclei

**DOI:** 10.1101/2020.07.27.222380

**Authors:** Salim Abdisalaam, Shibani Mukherjee, Souparno Bhattacharya, Debapriya Sinha, Sharda Kumari, Hesham A. Sadek, Janice Ortega, Guo-Min Li, Aroumougame Asaithamby

**Affiliations:** Department of Radiation Oncology, University of Texas Southwestern Medical Center, Dallas, Texas 75390.; Department of Internal Medicine, University of Texas Southwestern Medical Center, Dallas, Texas 75390.

**Keywords:** NBS1, cGAS, CtIP, dysfunctional telomeres, DNA damage response, micronuclei

## Abstract

Cyclic GMP-AMP synthase (cGAS), an important component of immune signaling, is hyperactivated in cells defective for DNA damage response (DDR) signaling. However, a direct role for DDR factors in the regulation of cGAS functions is mostly unknown. Here, we provide novel evidence that Nijmegen breakage syndrome 1 (NBS1) protein, a well-studied DNA double-strand break (DSB) sensor, in coordination with ATM, a protein kinase, and CtBP-interacting protein (CtIP), a DNA end resection factor, functions as an upstream regulator of cGAS binding to micronuclei. Upon NBS1 binding to micronuclei via its fork-head–associated domain, it recruits ATM and CtIP via its N- and C-terminal domains, respectively. Subsequently, ATM stabilizes NBS1’s interaction with micronuclei, and CtIP converts DSB ends into single-strand DNA ends, and these two key events preclude cGAS from binding to micronuclei. Notably, we show that purified cGAS cannot form a complex with DNA substrates that mimic resected DNA ends in vitro. Thus, NBS1 together with its binding partners modify the chromatin architecture of the micronuclei and that plays a critical role in cGAS’s binding to micronuclei.

## Introduction

Cyclic GMP-AMP synthase (cGAS), one of the most essential components of the innate immune system, is a cytosolic DNA sensor that is activated by binding to double-stranded DNA (dsDNA), including microbial and self-DNA. Upon binding to dsDNA in the cytoplasm, cGAS synthesizes 2’-3’-cGAMP, an atypical cyclic di-nucleotide second messenger that binds and activates STING (stimulator of interferon genes). STING then recruits TBK1 and IkB to activate IRF3 and NF-κB, respectively, leading to the production of type I interferons and inflammatory cytokines.^1–5^ A well-established function of cGAS in response to infections is to recognize microbial DNA linked to the orchestration of host defense programs.^2^ Through sensing self-DNA, such as the DNA of dead tumor cells, micronuclei, cytoplasmic chromatin fragments, and free telomeric DNA, cGAS engages the cGAS-STING pathway on both sides of cancer. On one side, this pathway is instrumental in antitumor immunity and promoting cellular senescence, intrinsic barriers in tumor cells that block malignancy. On the other side, the inflammatory consequences of the cGAS-STING pathway can also become maladaptive and promote inflammation-driven carcinogenesis and metastasis.^6–14^ cGAS is activated in response to genotoxic stress due to genomic instability or improper DNA damage response (DDR) signaling,^15^ but a relationship between DDR factors and the regulation of cGAS activation has not yet been established.

Nibrin or Nijmegen breakage syndrome protein 1 (NBS1) is a multifunctional protein that forms part of the MRN complex (MRE11–RAD50–NBS1) and is one of the key DDR factors. NBS1 is the protein defective in Nijmegen breakage syndrome (NBS), a rare autosomal recessive disorder associated with immune deficiency, microcephaly, chromosomal instability, and a high frequency of multiple malignancies.^16^ NBS1 participates in the early sensing of DNA damage and functions as a platform for recruiting and assembling the DDR components required for the repair process.^17^ NBS1 directly interacts with Meiotic Recombination 11 (MRE11),^18^ Ataxia Telangiectasia Mutated (ATM),^19,20^ Carboxy-terminal binding protein (CtBP)-interacting protein (CtIP),^21–23^ and the phosphorylated H2AX (γH2AX), a surrogate marker for DNA double-strand breaks (DSB). These NBS1 interacting proteins play key functions during different stages of DDR signaling. For example, NBS1-dependent recruitment of ATM to DSB sites not only phosphorylates NBS1 but also activates CtIP.^22^ CtIP binds to the N-terminal FHA-BRCT1/2 domains of NBS1; this binding is important for DNA end resection.^22,24^ Although NBS1 has been implicated in regulating inflammatory signaling,^25–28^ the contribution of NBS1 and its binding partners to this inflammatory signaling has not yet been identified.

In this study, we have discovered that NBS1’s DNA damage sensing characteristics play a key function in regulating cGAS’s ability to bind micronuclei in response to DNA damage. We found that NBS1 not only binds tightly to micronuclei harboring DSBs, but also recruits ATM and CtIP. Subsequently, ATM stabilizes NBS1 at the damaged chromatin, and CtIP converts DSB ends into single-strand DNA (ssDNA) ends. These ssDNA ends, and possibly the modification of chromatin organization during DNA end resection, block cGAS binding to all micronuclei. As the repair of DNA lesions within the micronuclei progresses and/or NBS1 binding to damaged chromatin weakens by unknown mechanisms, cGAS is able to interact with a limited number of micronuclei. Consequently, this limited cGAS binding activates innate immune signaling, which leads to cellular senescence in response to genotoxic stress. Thus, NBS1 constitutes an upstream regulatory factor for cGAS-mediated immune signaling and associated functions, including cellular senescence in response to DNA damage.

## Results

### cGAS is recruited to only a sub-set of micronuclei

A number of studies have reported the accumulation of cGAS in micronuclei in response to DNA damage and due to improper DDR signaling^14,29–32^; however, none have provided information on whether cGAS binds to all or to only a sub-set of micronuclei in response to DNA damage, and the role of DDR factors in regulating cGAS access to micronuclei. Initially, we evaluated the efficiency of cGAS recruitment to micronuclei in human cells in response to different types of DNA damaging agents, such as ionizing radiation (a genome-wide DSBs generator), aphidicolin, camptothecin, hydroxyurea and gemcitabine (replication stress inducers), and 6-thio-2’-Dideoxyguanine (6-thio-dG), dominant negative telomeric-repeat binding factor 2 (DN-TRF2), and CRISPR/Cas9 (sgTelomere-DD-Cas9; inducers of dysfunctional telomeres).^33^ Under similar number of micronuclei generation conditions (**Fig. 1A**), we observed that only a sub-set (~30-40%) of micronuclei were positive for cGAS, regardless of how we caused genotoxic stress (**Fig. 1B**). Among these DNA damaging agents, we noticed higher number of cGAS-positive micronuclei in response to dysfunctional telomeres caused by DN-TRF2, 6-thio-dG and sgTel-DD-Cas9. Additionally, we have recently shown that cGAS-STING-IRF3-activation in response to DN-TRF2, 6-thio-dG and sgTel-DD-Cas9 leads to cellular senescence.^33^ Therefore, we have chosen these dysfunctional telomere inducing strategies for uncovering DDR factors’ involvement in modulating cGAS binding to micronuclei.

**Figure 1:**
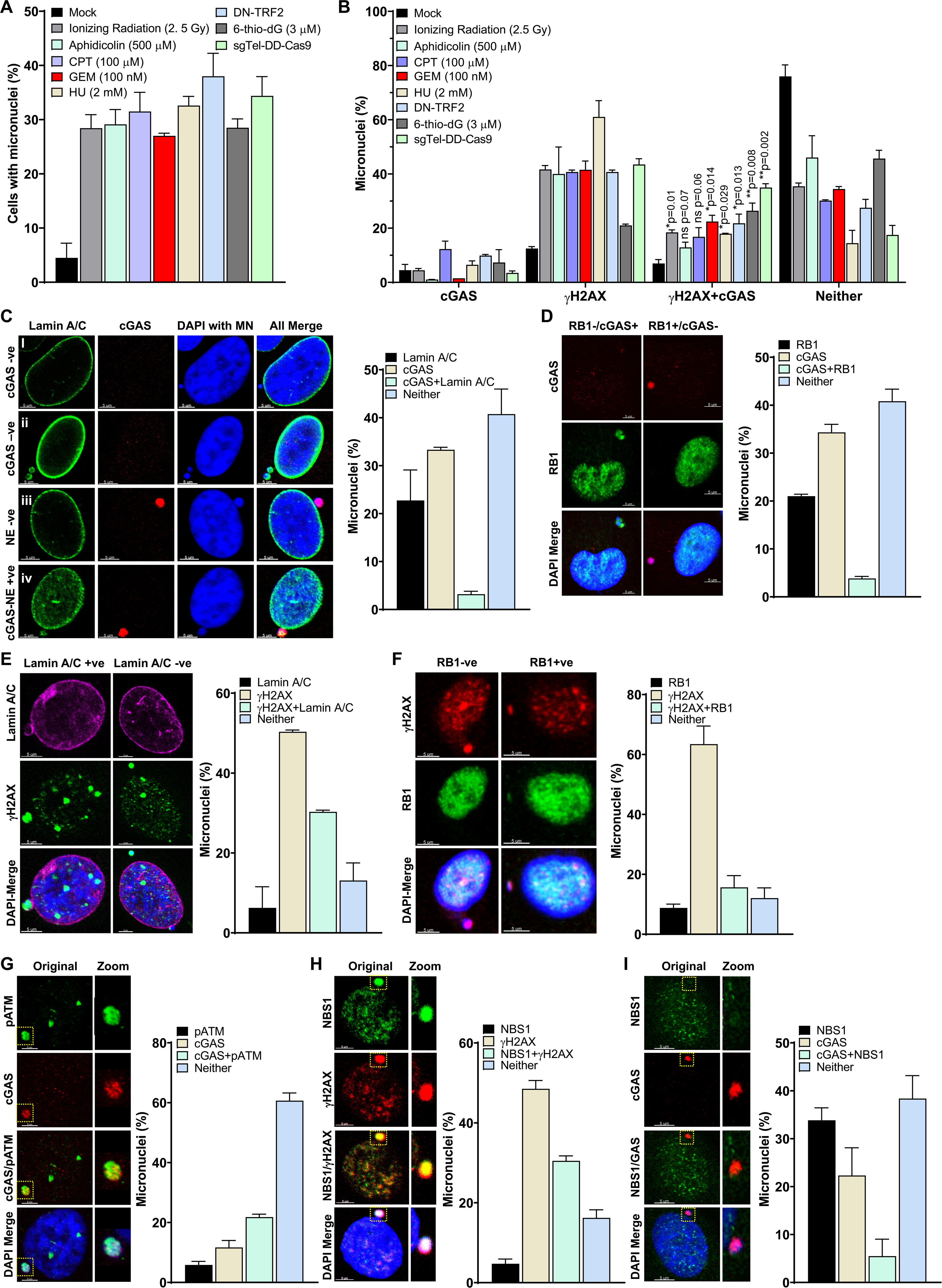
cGAS is recruited to only a sub-population of micronuclei in response to DNA damage. **A-B.** Only a limited number of micronuclei recruit cGAS in response to DNA damage. Bar graphs show percentages of micronuclei (**A**) and the percentage of micronuclei harboring either cGAS alone, γH2AX alone, cGAS and γH2AX or neither (**B**) at 24-72 hours after exposure of human bronchial epithelial (BEAS2B-APH/CPT/GEM/6-thio-dG) and HT1080 (DN-TRF2/sgTel-DD-Cas9) cells to different genotoxic agents. The bar graph presents the mean and STDEV from 400-600 cells from two independent experiments. Statistical analysis was performed using student’s t-test. APH-aphidicolin; CPT-camptothecin; GEM-gemcitabine; HU-Hydroxyurea; 6-thio-dG-6-thio-2’-deoxyguanine; DN-TRF2-overexpression of doxycycline inducible dominant negative telomeric-repeat binding factor 2; sgTel-DD-Cas9-CRISPR/Cas9 mediated induction of DNA double strand breaks in the telomeric-repeat DNA. **C-D.** Not all micronuclei (MN) with ruptured nuclear envelopes (NE) recruit cGAS. Representative images show cGAS- and Lamin A/C-negative (**i**), Lamin A/C-positive and cGAS-negative (**ii**), cGAS-positive and Lamin A/C-negative (**iii**) and cGAS-positive and Lamin A/C (weak) positive (**iv**) micronuclei (**C, left panels**). Bar graph shows the frequency of micronuclei harboring either Lamin A/C coating, cGAS, Lamin A/C and cGAS, or neither in BEAS2B cells treated with 3 μM 6-thio-dG for 72 hours (**C, right**). Representative images show localization of either cGAS or RB1 in micronuclei (**D, left**). Bar graph shows the frequency of micronuclei harboring either cGAS, RB1, cGAS and RB1 or neither in BEAS2B cells treated with 3 μM 6-thio-dG (**D, right**). Bar graph presents the mean and STDEV from 150-200 cells from two independent experimental groups. **E.** A major fraction of γH2AX-positive micronuclei are devoid of a nuclear envelope. Representative images show the presence or absence of γH2AX in Lamin A/C coating-positive and negative micronuclei (**left**). Bar graph shows the frequency of micronuclei harboring either Lamin A/C coating, γH2AX, Lamin A/C and γH2AX, or neither in BEAS2B cells treated with 3 μM 6-thio-dG for 72 hours (**right**). Bar graph presents the mean and STDEV from 50 different fields from two independent experimental groups. +ve-positive; −ve-negative. **F.** Representative images show the presence or absence of γH2AX in RB1-positive and RB1-negative micronuclei (**left**). Bar graph shows the percentage of micronuclei harboring either γH2AX, RB1, γH2AX and RB1 or neither in BEAS2B cells treated with 3 μM 6-thio-dG for 72 hours (**right**). Bar graph presents the mean and STDEV from 50 different fields from two independent experimental groups. **G.** Phosphorylated ATM (pATM, S1981) is recruited to micronuclei, and a fraction of it co-localizes with cGAS. Representative images show co-localization of pATM and cGAS in micronuclei (**left**). Bar graphs show the frequency of micronuclei containing pATM, cGAS, both or neither in BEAS2B cells treated with 3 μM 6-thio-dG (**right**). Bar graph presents the mean and STDEV from 50 different fields. **H-I.** NBS1 is recruited to γH2AX positive micronuclei but rarely co-localize with cGAS. Representative images show the presence of NBS1 and γH2AX (**H, left**) and only cGAS but no NBS1 (**I, left**) in micronuclei. Bar graphs show the frequency of micronuclei harboring NBS1, γH2AX, both or neither (**H, right**) and NBS1, cGAS, both or neither (**I, right**) in BEAS2B cells treated with 3 μM 6-thio-dG for 72 hours. Bar graph presents the mean and STDEV from 50 different fields.

Evidence indicates that, after micronuclei are formed, they are initially coated with a nuclear membrane, and rupturing this nuclear membrane exposes chromatin to the cytosolic environment. This phenomenon is required to recruit cGAS to the cytosolic chromatin fragments (CCFs) and to activate cytosolic DNA sensing pathway. Therefore, we examined the relationship between cGAS recruitment and nuclear envelope rupture by co-immunostaining after allowing cells to accumulate micronuclei in response to dysfunctional telomeres caused by 6-thio-dG. Similar to previous reports,^14,29–34^ we observed that most cGAS-bound micronuclei did not have a nuclear envelope coating (Lamin) (**Fig. 1C**). Interestingly, we noticed another population (~40%) of micronuclei that was neither Lamin- nor cGAS-positive (**Fig. 1C**). We further verified the limited recruitment of cGAS to micronuclei by co-immunostaining with RB1, another marker for intact nuclear envelope coating (**Fig. 1D**).^14,30,31^ As with previous reports, a major fraction of cGAS-positive micronuclei was negative for RB1, and some micronuclei were neither cGAS- nor RB1-positive (**Fig. 1D**). These observations raise the questions of why some micronuclei without any nuclear envelope coating do not recruit cGAS and whether factors those interact with micronuclei play any role in limiting cGAS recruitment to all micronuclei.

Apart from cGAS, we also detected the γH2AX signal, a surrogate marker for DSBs in a major fraction (up to 85%) of micronuclei (**Fig. 1B**), which suggests the presence of DSBs in these micronuclei. Though most of the cGAS-positive micronuclei were positive for γH2AX foci, not all γH2AX foci-positive micronuclei recruited cGAS (**Fig. 1B**). Furthermore, the micronuclei negative for Lamin coating (**Fig. 1E**) and RB1 (**Fig. 1F**) were also positive for γH2AX foci. Since several DNA damage sensing, repair and signaling factors, including ATM, MDC1, NBS1 (component of MRN complex), BRCA1, P53 and RPA32, are shown to localized in the micronuclei,^35,36^ it is reasonable to speculate that one of these factors may modulate the recruitment of cGAS to micronuclei in response to DNA damage. Initially, we examined the localization of MDC1, ATM, MRE11, and NBS1 to the micronuclei and found that ~18-40% of micronuclei were positive for MDC1, phosphorylated ATM (S1981), MRE11 and NBS1 in response to 6-thio-dG (**Figs. 1G-I**). We noticed co-localization of MDC1 (**Fig. S1A**) and ATM (**Fig. 1G**) with cGAS in the micronuclei. In contrast, we rarely detected co-localization of MRE11 (**Fig. S1B**) and NBS1 (**Figs. 1H-I**) with cGAS in the same micronuclei, which indicates a mutually exclusive relationship between NBS1/MRE11 and cGAS with regard to binding to micronuclei.

### NBS1 deficiency enhances cGAS recruitment to micronuclei

Our finding that only a minor fraction of NBS1 co-localizes with cGAS in micronuclei prompted us to investigate NBS1’s involvement in cGAS recruitment to these micronuclei in response to dysfunctional telomeres (6-thio-dG). First, we used shNBS1 RNA (Sigma) to stably knockdown NBS1 in BEAS2B cells (**Fig. 2A**). Surprisingly, although we observed a comparable number of micronuclei in scrambled shRNA (shSCR) and shNBS1 RNA cells (**Fig. 2B**), we noticed that ~66% of the micronuclei in shNBS1 RNA cells were positive for cGAS, but only 33% of the micronuclei in shSCR cells were positive for cGAS in response to 6-thio-dG (**Fig. 2C**). To rule out the possibility that the increase in cGAS-positive micronuclei is not associated with genomic instability due to long-term-knockdown of NBS1, we used a tetracycline-inducible shNBS1 RNA. As with the stable-knockdown of NBS1, we noticed significantly elevated levels of cGAS-positive micronuclei (~59%) in tetracycline-mediated NBS1-knockdown cells in response to 6- thio-dG (**Fig. 2C**). Significantly, these increased numbers of cGAS-positive micronuclei were associated with increased phosphorylation of IRF3 and STAT1 (**Fig. 2D**), elevated levels of IL-6 secretion (**Fig. 2E**), and higher expression levels of genes associated with immune signaling (**Fig. 2F**) in NBS1-knockdown cells relative to cells expressing control shRNA. Ultimately, we detected higher incidences of β-gal–positive stable and tetracycline-inducible NBS1-knockdown cells than in the control shRNA-expressing cells in response to 6-thio-dG (**Fig. 2G**).

**Figure 2:**
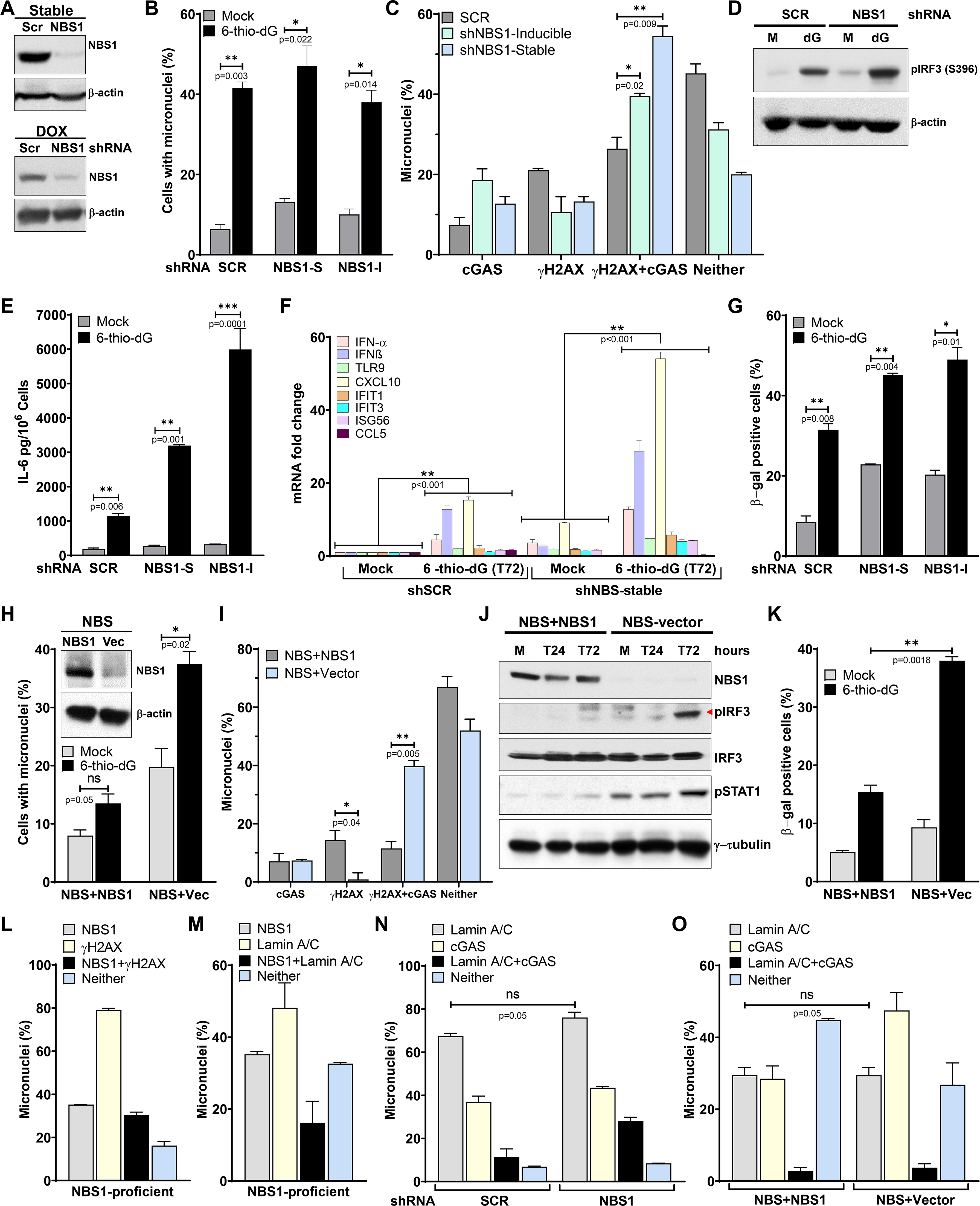
NBS1 deficiency enhances cGAS-positive micronuclei. **A-C.** Stable and transient NBS1 knockdown enhances cGAS-positive micronuclei. Western blots show stable (top) and doxycycline (DOX; lower) mediated NBS1 depletion in BEAS2B cells (**A**). Bar graph shows percentages of micronuclei in mock- and 3 μM 6-thio-dG (72 h) treated BEAS2B cells stably expressing scrambled (SCR), stable (NBS1-S), and doxycycline inducible (NBS1-I) shNBS1 RNAs (**B**). Bar graph shows the percentage of micronuclei harboring either cGAS alone, γH2AX alone, cGAS and γH2AX or neither in scrambled (SCR), stable, and doxycycline inducible shNBS1 RNA expressing BEAS2B cells at 72 hours after 3 μM thio-dG treatment (**C**). Bar graph presents the mean and STDEV from 400-600 cells from two independent experiments and the statistical analysis was performed using student’s t-test. **D-G.** NBS1 knockdown enhances IRF3 phosphorylation, IL-6 secretion, expression of immune gene and cellular senescence. Sample western blot shows phosphorylation of IRF3 (S396; **D**), and the bar graph shows the level of IL-6 in the BEAS2B cell culture supernatant (**E**) in mock- and at 72 hours after 3 μM 6-thio-dG (dG) treatment. Graph shows greater expression of immune pathway genes in 3 μM 6-thio-dG treated shNBS1 RNA cells than in shSCR RNA cells (**F**). Bar graph shows frequency of β-gal staining-positive cells at 10 days after 3 μM 6-thio-dG treatment in shSCR, shNBS1-stable (S) and doxycycline inducible (I) RNA-expressing BEAS2B cells (**G**). Bar graph presents the mean and STDEV from 10-15 fields from two independent experiments. Statistical analysis was performed using student’s t-test (E and G) and two-way ANOVA (F). **H-I.** Elevated levels of cGAS-positive micronuclei in patient-derived NBS1-mutant (NBS) cells. Western blots show expression of NBS1 in NBS complemented with NBS1 and vector-alone (vec) cells (**H, inset**). Bar graph shows the percentage of micronuclei in mock- and 5 μM 6-thio-dG (72 h) treated NBS1-mutant (NBS+Vec) and NBS cells complemented with WT NBS1(NBS+NBS1; **H**). Bar graph shows the percentage of micronuclei harboring either cGAS alone, γH2AX alone, cGAS and γH2AX or neither in NBS1-mutant (NBS+Vector) and NBS cells complemented with WT NBS1 (NBS+NBS1) at 72 hours after 5 µM 6-thio-dG treatment. Bar graphs present the mean and STDEV from 50 different fields from two independent experiments. Statistical analysis was performed using student’s t-test. **J-K.** NBS1 deficiency exacerbates premature senescence. Western blots show increased activation of IRF3 (S396) and STAT1 (Y701) at different times after 6-thio-dG treatment (**J**). NBS cells show higher levels of β-gal staining than NBS+NBS1 cells 7 days after 5 μM 6-thio-dG treatment (**K**). Bar graph presents the mean and STDEV from 12-15 different fields from two independent experiments. Statistical analysis was performed using student’s t-test. **L.** A major fraction of NBS1 co-localizes with DSB marker, γH2AX. Bar graph shows the frequency of micronuclei containing either DSB marker (γH2AX), NBS1, both or neither in 3 μM 6-thio-dG-treated BEAS2B cells. Data in the bar graph present the mean and STDEV from 50 different fields from two independent groups. **M.** NBS1 recruitment to micronuclei is independent of nuclear envelope coating. Bar graph shows the frequency of micronuclei containing either nuclear envelope marker (Lamin A/C) alone, NBS1 alone, both or neither in 3 μM 6-thio-dG-treated BEAS2B cells. Data in the bar graph present the mean and STDEV from 50 different fields from two independent groups. **N-O.** Increased proportion of cGAS-positive micronuclei in NBS1-knockdown (**N**) and NBS (**O**) cells is not due to micronuclei’s defective nuclear envelope coating. Bar graphs show the percentage of micronuclei harboring either Lamin A/C coating, cGAS, Lamin A/C+cGAS or neither in SCR- and doxycycline inducible shNBS1 RNAs-expressing BEAS2B (**N**) and in NBS cells stably expressing either vector alone or FL-NBS1 (**O**) 72 hours after 6-thio-dG (3-5 µM) treatment. Bar graphs present the mean and STDEV from 50 different fields from two independent experiments. Statistical analysis was performed using student’s t-test.

We validated these results by using patient-derived NBS1-deficient NBS cells and NBS cells complemented with wild-type NBS1 (NBS+WT NBS1).^37^ Initially, we noticed a significantly higher frequency of micronuclei in NBS cells than in NBS+WT NBS1 cells in response to 6-thio-dG (**Fig. 2H**). Subsequently, we found that a significantly higher percentage of micronuclei had cGAS in cells lacking NBS1 than in NBS+WT NBS1 cells (**Fig. 2I**). Furthermore, IRF3 activation, as assessed by western blotting (**Fig. 2J**) was markedly higher in NBS-deficient cells than in NBS+WT-NBS1 cells in response to 6- thio-dG. Additionally, NBS1-deficient cells exhibited elevated levels of senescence relative to NBS+WT NBS1 cells in response to 6-thio-dG (**Fig. 2K**). Thus, NBS1 is somehow involved in negatively regulating cGAS binding to micronuclei.

Because NBS1 is known to be recruited to the sites of DSBs, we evaluated whether all NBS1 co-localizes with γH2AX-positive micronuclei or is independent of DSBs in the micronuclei. We found that a major percentage (80-90%) of NBS1-positive micronuclei harbor γH2AX signals in BEAS2B cells treated with 6-thio-dG, which implies that NBS1 is localized to DSB-positive micronuclei (**Fig. 2L**). Additionally, we evaluated whether NBS1 localization is restricted to micronuclei whose nuclear envelope is intact or ruptured. Co-immunostaining BEAS2B cells with NBS1 and Lamin A/C antibodies revealed that 35-40% of NBS1-positive micronuclei are also positive for Lamin A/C, and the rest are devoid of any Lamin A/C staining (**Fig. 2M**). We corroborated these results by co-staining BEAS2B cells with NBS1 and RB1 antibodies. Like Lamin A/C-NBS1 co-localization, only 15-20% micronuclei are positive for RB1 and NBS1, and the remaining NBS1-positive micronuclei are devoid of RB1 signals (**data not shown**). Thus, localization of NBS1 in micronuclei is dependent on γH2AX but independent of nuclear envelope coating.

Subsequently, to rule out the possibility that the increased number of cGAS-positive micronuclei resulted from a defective nuclear envelope coating, we counted the nuclear envelope-positive and negative micronuclei in shSCR RNA, shNBS1 RNA, NBS and NBS+WT NBS1 cells. We found no statistically significant difference in the number of nuclear envelope-coated micronuclei between shSCR and shNBS1 (**Fig. 2N**), nor between NBS and NBS+WT NBS1 cells (**Fig. 2O**). Therefore, the increased percentage of cGAS-positive micronuclei in NBS1-knockdown and NBS cells is not due to defects in the nuclear envelope coating, but to NBS1 deficiency.

### The FHA-domain–dependent recruitment of NBS1 to DSB sites and the CtIP binding domain of NBS1 are critical for attenuating cGAS binding to micronuclei

To narrow down the NBS1 domain that is responsible for modulating cGAS binding to micronuclei, we used NBS cells stably expressing a panel of NBS1 mutants, including full-length (FL)-NBS1 harboring two mis-sense point mutations (Gly 27 Asp and Arg 28 Asp; FHA-2D) in the FHA domain, and FHA-(ΔFHA), ATM recruitment domain-(Δ703- 754 or ΔATM) and MRE11 binding domain-(ΔMRE11; 682–693 amino acids) deleted NBS1 (**Figs. 3A**). These NBS1 mutants prevent recruitment to DSB sites of either NBS1 (FHA-2D and ΔFHA) or its binding partners, specifically CtIP (ΔFHA), ATM (ΔATM) and MRE11 (ΔMRE11).^16–20,38^ Initially, we noticed that the number of micronuclei induced by 6-thio-dG was higher in these NBS1 mutant cells than in NBS+WT NBS1 cells (**Fig. 3B**). Interestingly, we found that the levels of cGAS-positive micronuclei were significantly higher in NBS cells expressing FHA-2D-, ΔFHA- and ΔATM-NBS1 than in NBS cells expressing WT-NBS1 and ΔMRE11-NBS1 (**Fig. 3C**). Thus, the FHA-domain– dependent interaction of NBS1 with the DSB sites, and the CtIP and the ATM binding domains, but not the MRE11-binding domain, of NBS1 may be involved in modulating cGAS recruitment to those micronuclei.

**Figure 3.**
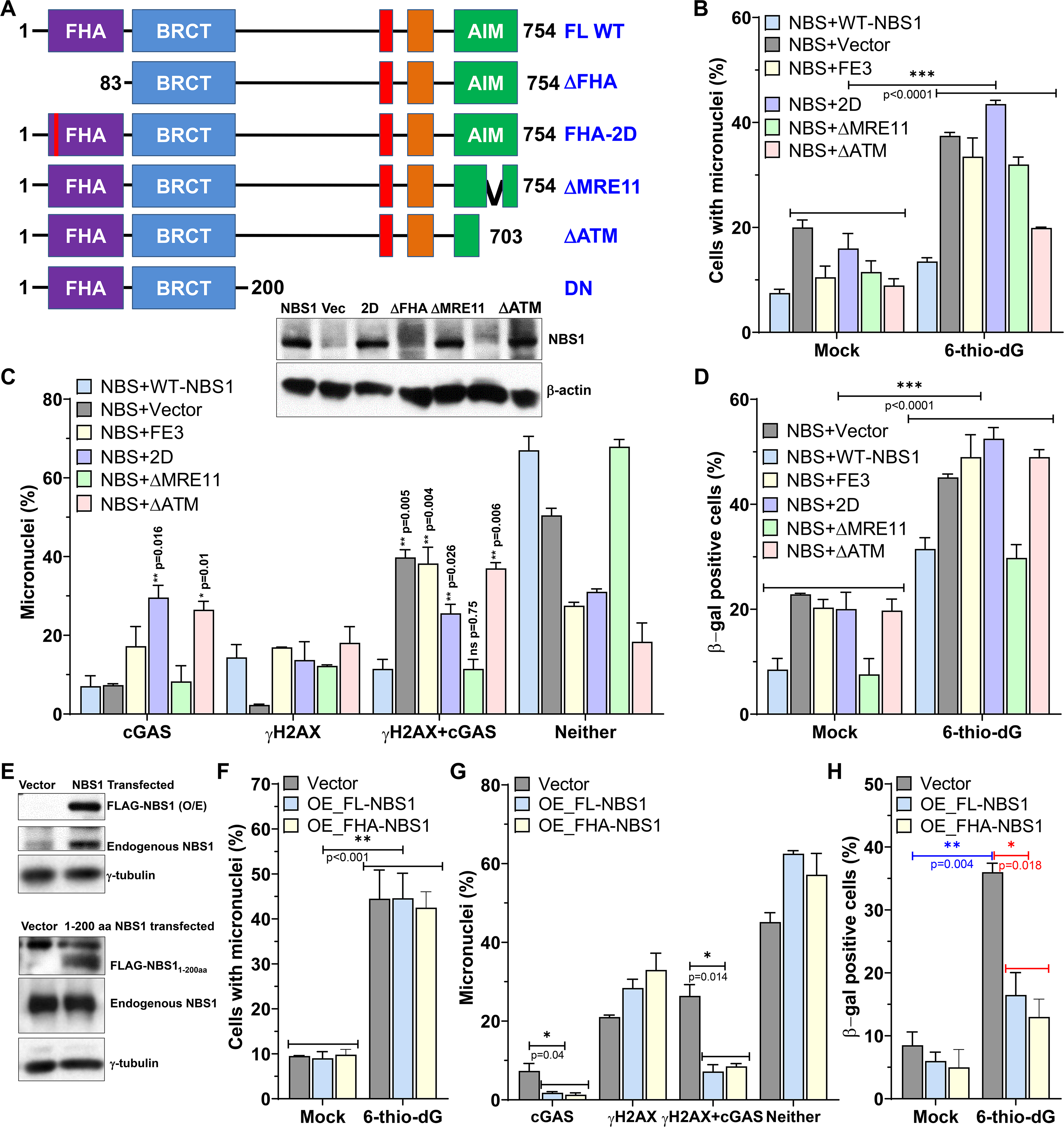
Physical presence of NBS1 at the DSB sites of micronuclei is critical for modulating cGAS binding to micronuclei. **A.** Schematics of different NBS1 mutant constructs used in this study (top). Western blot shows stable expression of different NBS1 mutants in NBS patient derived cells (lower). FHA-fork head-associated domain; BRCT-BRCA1/2 C-terminus domain; AIM-ATM-MRE11 interaction domain; DN-dominant negative NBS1. **B-C.** Defects in the FHA and ATM binding domains of NBS1 causes increased numbers of cGAS positive micronuclei and senescent cells. Bar graph shows percentages of micronuclei in NBS cells stably expressing FHA-2D, ΔFHA, ΔMRE11 and ΔATM NBS1 mutants at 72 hours after mock and 5 μM thio-dG treatment (**B**). Bar graph shows the percentage of micronuclei harboring either cGAS alone, γH2AX alone, cGAS and γH2AX or neither in NBS cells stably expressing FHA-2D, ΔFHA, ΔMRE11 and ΔATM NBS1 mutants at 72 hours after 5 μM thio-dG treatment (**C**). NBS cells stably expressing NBS-2D, ΔFHA and ΔATM NBS1 mutants show higher levels of β-gal staining than NBS+NBS1 cells 7 days after 5 μM 6-thio-dG treatment (**D**). NBS+Vector and NBS1+FL-NBS1 data from figure 2 was used for comparison. Bar graph presents the mean and STDEV from 400-600 cells from two independent experiments. Statistical analysis was performed using two-way ANOVA (B and D) and student’s t-test (C). **E-H.** Physical presence of NBS1 within the micronuclei is required to block cGAS recruitment to micronuclei. Overexpression (O/E) of full-length (FL) and FHA-BRCA domain (NBS1_1-200aa_): i.e. dominant negative NBS1-DN) NBS1 in NBS1-proficient BEAS2B cells reduces cGAS-positive micronuclei. Western blot shows O/E of FL (top) and NBS1_1-200aa_ (lower) NBS1 in BEAS2B cells (**E**). Graph shows the percentages of micronuclei in BEAS2B-cells overexpressing FL and NBS1_1-200aa_ (DN) NBS1 in mock and at 72 hours after 3 μM 6-thio-dG treatment (**F**). Bar graph shows the percentage of micronuclei harboring either cGAS alone, γH2AX alone, cGAS and γH2AX or neither in BEAS2B-cells overexpressing FL and NBS1_1-200aa_ (DN) NBS1 at 72 hours after 3 μM 6-thio-dG treatment (**G**). Bar graph shows the percentage of β-gal staining positive BEAS2B-cells O/E FL and NBS1_1-200aa_ (DN) NBS1 at 7 days after 3 μM 6-thio-dG treatment (**H**). Bar graphs present the mean and STDEV of two-three independent experiments. Statistical analysis was performed using two-way ANOVA (F) and student’s t-test (G and H).

To validate that the physical interaction of NBS1 with the DSB-harboring micronuclei is necessary to modulate cGAS binding to micronuclei, we stably overexpressed full-length (FL) and 1–200 amino acid region-containing FHA and BRCT1 domains (NBS1_1-200aa_ or DN-NBS1) of NBS1 in NBS1-proficient cells (**Fig. 3D**) and examined cGAS recruitment to micronuclei in response to 6-thio-dG. FL-NBS1 and DN-NBS1 overexpression will lead to enhanced binding of FL- and DN-NBS1 and its binding partners to micronuclei. We noticed significantly reduced numbers of cGAS-positive micronuclei (**Fig. 3E**), but not numbers of micronuclei (**Fig. 3F**), in cells overexpressing FL-NBS1 and NBS1_1-200aa_ in response to 6-thio-dG. Thus, the physical association of NBS1 with the DSB sites of the micronuclei somehow limits cGAS recruitment to those micronuclei. Finally, we found that neither MRE11 nor its exonuclease activity is involved in the regulation of cGAS recruitment to micronuclei in response to 6-thio-dG (**Fig. S2** and **supplementary results**).

### NBS1, but not cGAS, occupies chromatin fragments in metaphase, and its chromatin binding affinity decreases during metaphase to telophase transition

Thus far, we have confirmed that cGAS is recruited to only a limited population of micronuclei in NBS1-proficient cells and its binding efficiency to micronuclei is significantly enhanced in NBS1-defective cells in response to genotoxic stress. However, how cGAS binds to certain micronuclei and why some micronuclei do not recruit cGAS are still unclear. First, we set to determine how cGAS binds to some micronuclei in NBS1-proficient cells in response to genotoxic stress. Multiple mechanisms are possible, including the timing of NBS1 binding to damaged chromosome fragments and the difference in dynamic interactions between NBS1 and cGAS on the damaged chromosome fragments. Reports suggest that NBS1 can bind to damaged chromatin in the metaphase stage,^39^ whereas cGAS is recruited to the chromatin fragments during telophase stage.^30^ To test these previous observations, we performed live cell imaging in cells co-expressing TagRFP-cGAS and EGFP-NBS1. We observed that EGFP-NBS1, but not TagRFP-cGAS, bound to mis-aligned chromosome in metaphase cells (**Fig. 4A**). Additionally, we found that NBS1 remained anchored to mis-segregated chromosome when cells progressed to the anaphase stage of mitosis (**Fig. 4A**). Interestingly, we noticed a gradual accumulation of TagRFP-cGAS onto some of these chromatin fragments during early telophase, and the cGAS fluorescence signal increased during the transition to late telophase (**Fig. 4A**).

**Figure 4:**
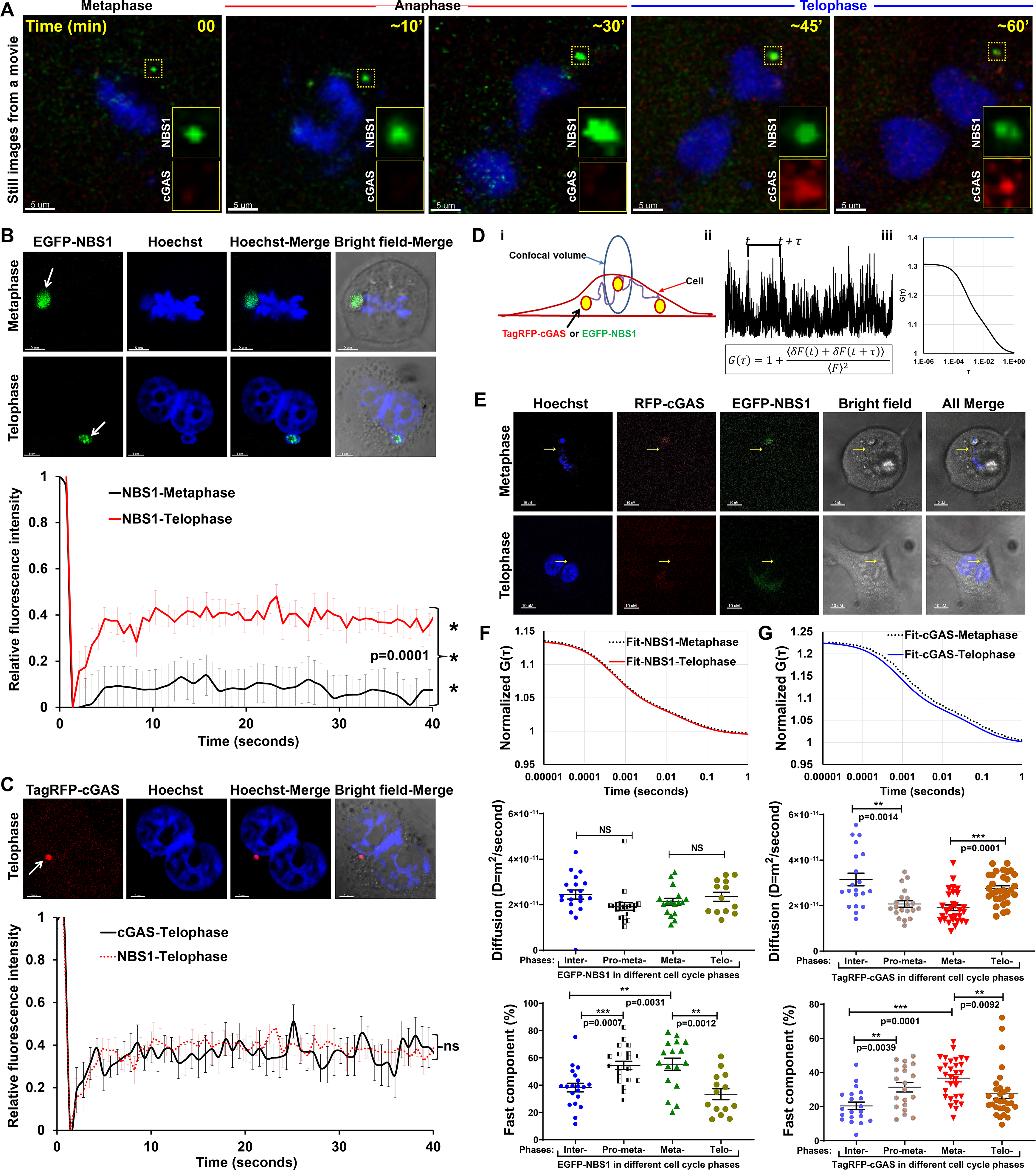
NBS1 is recruited during metaphase through telophase, and it binds to metaphase chromatin more tightly than cGAS. **A.** NBS1 is recruited to mis-segregated chromatin fragments in metaphase whereas cGAS is recruited in telophase. Representative still images from a movie show recruitment and retention of EGFP-NBS1 in metaphase and telophase, respectively, and recruitment of Tag-RFP-cGAS in telophase in BEAS2B cells co-expressing EGFP-NBS1 and doxycycline-inducible Tag-RFP-cGAS. EGFP-NBS1 and doxycycline-inducible Tag-RFP-cGAS cells were co-treated with 3 µM 6-thio-dG and 250 ng/ml doxycycline for 72 hours, mitotic cells were collected by mitotic shake-off and were plated onto a glass-bottomed cell culture dish. 2-4 hours after seeding the cells, they were subjected to live cell imaging for 8-12 hours using LSM510 Meta (Zeiss) microscope equipped with CO_2_ and temperature-controlled cell culture chambers. **B. C.** NBS1’s binding affinity to chromatin fragments in metaphase is tighter than telophase. Representative live cell images show EGFP-NBS1 fluorescence recovery after photobleaching (FRAP) regions (white arrows) of chromatin fragments in metaphase and telophase cells (**B, top**). Line graphs show FRAP curves of EGFP-NBS1 in metaphase and telophase (**B, bottom**). Representative live cell images show Tag-RFP-cGAS FRAP region of chromatin fragment in telophase (**C, top**) and the line graph show FRAP curves of Tag-RFP-cGAS in telophase (**C, lower**) in BEAS2B cells stably expressing EGFP-NBS1 and doxycycline-inducible Tag-RFP-cGAS treated with 3 μM 6-thio-dG. EGFP-NBS1 FRAP curve from panel **B** was used for comparison. Each data point depicted in the graph is the mean and STDEV from 13-30 measurements taken in 13-30 different cells at different pre- and post-FRAP times. Data represent three independent experiments. Statistical analysis was performed using two-way ANOVA. **D.** Diagram of FCS methodology, showing how temporal analysis of fluorescence fluctuations (**i**) is converted into the autocorrelation function (ACF; **ii**), which is related to diffusion time (**iii**). **E-G.** Representative live cell images show EGFP-NBS1 and Tag-RFP-cGAS fluorescence correlation spectroscopy (FCS) regions (yellow arrows) adjacent to the chromatin fragments in metaphase and telophase cells (**E**). Representative FCS curve fits of EGFP-NBS1 in metaphase (dotted line) and in telophase (solid line; **F, top**); and TagRFP-cGAS in metaphase (dotted line) and telophase (solid line; **G, top**). Dot blots show diffusion coefficient values for EGFP-NBS1 (**F, middle**) and Tag-RFP-cGAS (**G, middle**); and the percentage of fast components of EGFP-NBS1 (**F, lower**) and Tag-RFP-cGAS (**G, lower**) in indicated cell cycle stages of BEAS2B cells stably co-expressing EGFP-NBS1 and doxycycline-inducible Tag-RFP-cGAS co-treated with 250 ng/ml doxycycline and 3 μM 6-thio-dG for 72 hours. Each point of the dot-plot represents measurement from a single cell produced from 20 x 5-s reading. Values show the mean from >20 measurements taken in 15-30 individual interphase (Inter-), pro-metaphase (pro-meta-), metaphase (Meta-) and telophase (Telo-) cells. Statistical analysis was performed using student’s t-test.

Why doesn’t NBS1 continue to bind to mis-segregated chromosomes in telophase, and what makes cGAS bind to mis-segregated chromosomes in telophase but not in metaphase? This could be due to NBS1’s and cGAS’s differential binding affinities with the mis-segregated chromosomes in different stages of the mitotic phase. First, we measured the dynamic exchange between free and mis-segregated chromosomes-bound EGFP-NBS1 in metaphase and telophase using the fluorescence recovery after photobleaching (FRAP) technique. Mis-segregated chromosomes-bound EGFP-NBS1 in metaphase recovered to only ~10% after photobleaching, whereas EGFP-NBS1 in telophase recovered to ~40% within 5 seconds after photobleaching (**Fig. 4B, lower panel**). These results imply that NBS1 binds more strongly to mis-segregated chromosomes in metaphase than to telophase chromosome fragments (p=0.0001). Similarly, chromosome fragments-bound TagRFP-cGAS in telophase recovered to ~40% within 5 seconds after photobleaching (**Fig. 4C, lower panel**). This observation indicates that there is a dynamic exchange between free floating cGAS and chromosome fragments-bound cGAS in telophase. Thus, NBS1 binds to chromosome fragments tightly in metaphase and that NBS1 and cGAS have comparable binding affinities for chromosome fragments in telophase.

Because of the relatively low abundance of cGAS and its lack of binding to chromatin fragments in metaphase, we were unable to analyze cGAS by FRAP in this mitotic stage. Instead, we measured the mobility (or diffusion) characteristics of cGAS molecules in different mitotic stages and compared them with those of NBS1 using fluorescence correlation spectroscopy (FCS), which is well suited for analyzing fast-diffusing proteins present in low concentrations. FCS records fluorescence intensity fluctuations in a small confocal volume as a function of time (**Fig. 4D**), allowing the rate (slow and fast) of diffusion properties of fluorescently tagged proteins within the complex and heterogeneous cellular environment to be determined. Utilizing an experimental setup similar to that described previously,^40^ we characterized the mobility of TagRFP-cGAS and EGFP-NBS1 near the vicinity of chromatin fragments (**Fig. 4E**, yellow arrows). We found that both NBS1 and cGAS molecules exist in complex states of mobility near the chromatin fragments (**Figs. 4F-G, top panels**). Their biological mobility behavior can be inferred as transient or stable complexes near or around the chromatin fragments. Specifically, we observed that, as mitotic phase cells transition from metaphase to telophase, cGAS mobility is significantly enhanced, as the diffusion coefficient changes from D=19±1.3 μm^2^/s in metaphase to D=27.5±1.2 μm^2^/s in telophase (p=0.0001; **Fig. 4G**, middle panel). Additionally, the percentage of fast components of cGAS reduced from ~36% (36 ±2.7%) in metaphase to ~27% (27 ±2.1%) in telophase (p=0.0092; **Fig. 4G**, lower panel). These results imply that the distribution of cGAS molecules changes towards a more stable steady state that is ready to interact with chromatin fragments and with other complexes around or on the chromatin fragments during telophase.

In contrast, we found that, as the mitotic phase progresses from metaphase to telophase, NBS1 mobility is slightly increased (but not significantly), as the diffusion coefficient changes from D=21.3±1.5 μm^2^/s in metaphase to D=23.5±2.0 μm^2^/s in telophase (**Fig. 4F**, middle panel). However, the percentage of fast components of NBS1 is significantly reduced from ~55% (55.5 ±4.5%) in metaphase to ~33% (33.3 ±4.0%) in telophase (p=0.0012; **Fig. 4F**, lower panel). These observations, together with FRAP results, indicate that NBS1’s tight binding affinity with the chromatin fragments and its slower diffusion characteristics may allow the steady-state localization of NBS1 in metaphase chromatin fragments, which may prevent cGAS from binding to chromatin fragments in metaphase. In telophase, by contrast, the changing NBS1 molecule distribution from chromatin-bound complexes to slow (free) components may provide an opportunity for cGAS molecules to access chromatin fragments.

### ATM-mediated NBS1 stabilization on chromatin fragments attenuates cGAS recruitment

Early recruitment, the dynamic interaction with chromatin fragments, and the diffusion properties are NBS1’s three major characteristics that modulate cGAS’s ability to interact with chromatin fragments. Therefore, factors that modulate the dynamic interaction between NBS1 and damaged chromatin can indirectly impact cGAS recruitment to chromatin fragments. NBS1 dynamics at damaged DNA are regulated by ATM-mediated phosphorylation.^41^ Evidence shows that NBS1 recruits ATM to DSB sites via its C-terminal regions (703–754 amino acids),^42^ resulting in NBS1 phosphorylation as well as NBS1 stabilization at DSB sites.^19,41^ Furthermore, ATM deficiency has been shown to enhance innate immune signaling.^43–46^ Additionally, we have demonstrated that the ATM-recruitment domain (Δ703-754 or ΔATM) of NBS1 is critical for attenuating cGAS binding to micronuclei in response to 6-thio-dG (**Figs. 3B-C**).

To evaluate ATM’s role in modulating cGAS binding to micronuclei, we initially examined the co-localization of NBS1 and ATM in micronuclei. We found that a major percentage of micronuclei harboring NBS1 was positive for phosphorylated ATM (S1981; **Fig. 5A**). Subsequently, we stably knocked-down ATM using ATM-specific shRNA. We found that shRNA-mediated ATM knockdown did not affect micronuclei formation in mock-treated cells, but ATM-knockdown cells showed higher numbers of micronuclei in response to 6-thio-dG (**Fig. 5B**). Furthermore, ATM--knockdown cells exhibited higher numbers of cGAS-positive micronuclei (**Fig. 5C**) and enhanced IRF3 activation (**Fig. 5D**) in response to 6-thio-dG.

**Figure 5:**
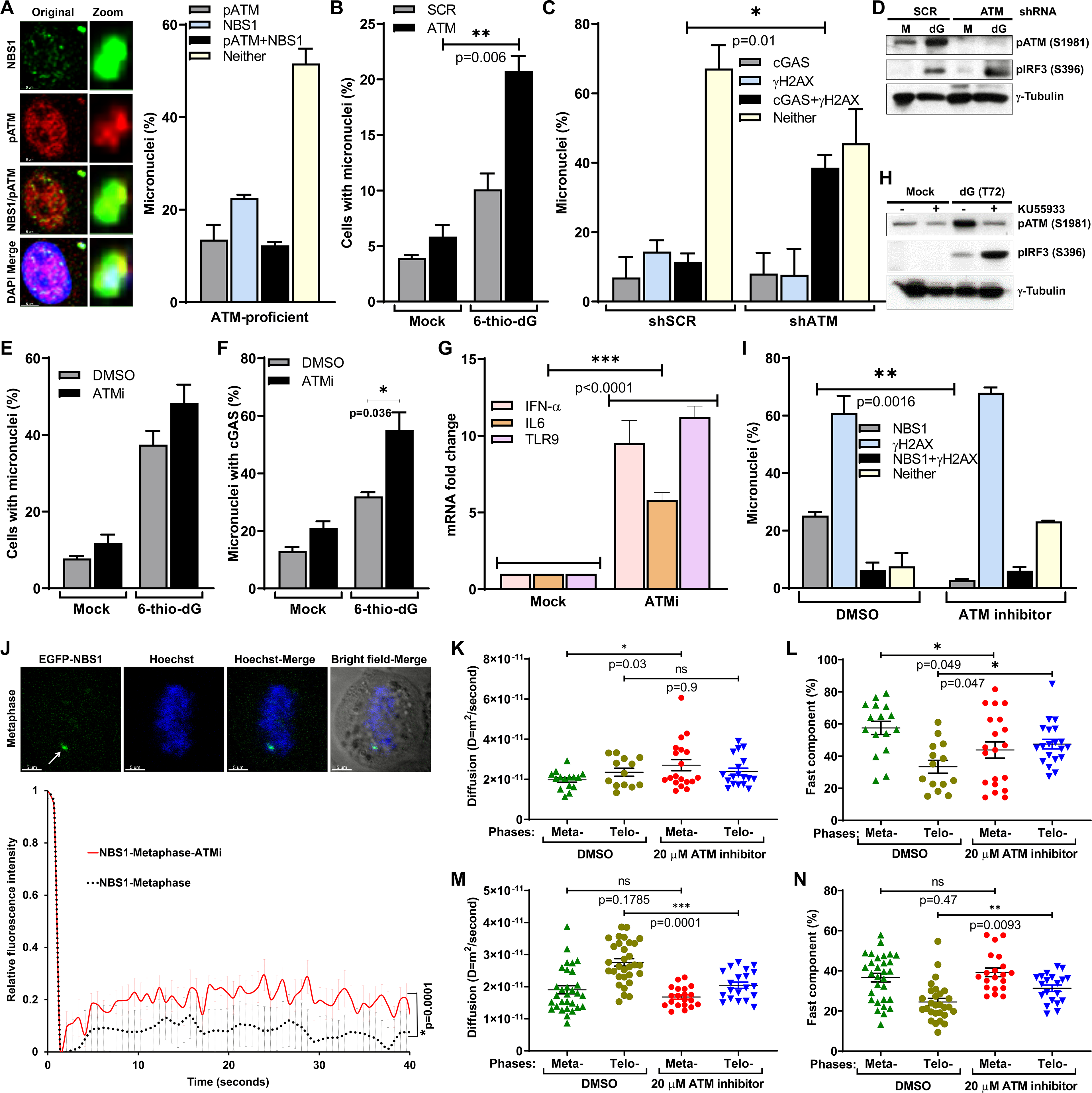
ATM-mediated NBS1 stabilization at DSB sites of micronuclei modulates cGAS binding to micronuclei. **A.** NBS1 co-localizes with ATM in micronuclei. Representative images show co-localization of NBS1 and phosphorylated ATM (S1981; **left**). Bar graph shows the frequency of micronuclei harboring either NBS1 alone, pATM alone, NBS1+pATM or neither in BEAS2B cells treated with 6-thio-dG (72 hours). Bar graph presents the mean and STDEV from 50 different fields (300-500 cells) from two independent experiments. **B-D.** ATM knockdown increases cGAS-positive micronuclei. Bar graph shows the frequency of micronuclei formation in mock- and 3 µM 6-thio-dG treated SCR and ATM shRNA cells (**B**). Bar graph show elevated levels of cGAS-positive micronuclei (**C**), and western blot shows increased levels of IRF3 phosphorylation (**D**) in shRNA-mediated ATM-knockdown cells than in SCR shRNA-expressing cells 72 hours after 3 µM 6-thio-dG treatment. Bar graph presents the mean and STDEV from 50 different fields of each experimental group. Statistical analysis was performed using student’s t-test. **E-H.** ATM kinase activity modulates cGAS recruitment to micronuclei. Bar graphs show the frequency of micronuclei formation (**E**), elevated levels of cGAS-positive micronuclei (**F**) and expression of IFNα, IL6 and TLR9 in cells treated with ATM kinase inhibitor (Ku55933) than the DMSO-treated cells exposed to 3 µM 6-thio-dG for 72 h. Bar graph presents the mean and STDEV from 50 different fields of each experimental group from two independent experiments. Statistical analysis was performed using student’s t-test. Western blots show elevated IRF3 phosphorylation in ATM inhibitor (Ku55933)-treated cells than in DMSO-treated BEAS2B cells exposed to 3 µM 6-thio-dG for 72 h (**H**). **I.** Inhibiting ATM kinase activity reduces NBS1 recruitment to micronuclei. Bar graph shows the percentage of micronuclei containing γH2AX, NBS1, both or neither in DMSO- and ATM inhibitor (Ku55933)-treated BEAS2B cells exposed to 3 µM 6-thio-dG for 72 h. Bar graph presents the mean and STDEV from 50 different fields of each experimental group from two independent experiments. Statistical analysis was performed using student’s t-test. **J.** Inhibiting ATM kinase activity reduces NBS1’s binding affinity to chromatin fragments in metaphase. Representative live cell images show EGFP-NBS1 FRAP region (white arrows) of chromatin fragments in metaphase. Line graph shows FRAP curves of EGFP-NBS1 in DMSO- and 20 μM ATM inhibitor (Ku55933)-treated BEAS2B cells co-expressing EGFP-NBS1 and doxycycline-inducible Tag-RFP-cGAS co-treated with 3 µM 6-thio-dG and doxycycline for 72 h. EGFP-NBS1 FRAP curve from figure 4B was used for comparison. Each data point depicted in the graph is the mean from >20 measurements taken in 20 different cells. Data represent three independent experiments. Statistical analysis was performed using two-way ANOVA. **K-N.** Dot blots show diffusion coefficient values for EGFP-NBS1 (**K**) and Tag-RFP-cGAS (**M**); and the percentage of fast components of EGFP-NBS1 (**L**) and Tag-RFP-cGAS (**N**) in metaphase and telophase BEAS2B cells stably co-expressing EGFP-NBS1 and doxycycline-inducible Tag-RFP-cGAS. Cells were first treated with 3 μM 6-thio-dG and doxycycline for 72 hours and then with either DMSO or 20 μM Ku55933 for 24 hours. Each point of the dot-plot represents measurement from a single cell produced from 20 x 5-s reading. Values show the mean from >20 measurements taken in 15-30 individual metaphase (Meta-) and telophase (Telo-) cells. Statistical analysis was performed using student’s t-test.

To determine whether ATM’s kinase activity is important for modulating cGAS binding to micronuclei, we inhibited this activity by using a small molecule inhibitor, Ku55933. Similar to ATM knockdown cells, we observed increased numbers of cGAS-positive micronuclei, expression of immune signaling genes and activation of IRF3 after inducing genotoxic stress with 6-thio-dG (**Figs. 5E-H**). Overall, ATM and its kinase activity are involved in regulating cGAS recruitment to micronuclei and subsequent activation of immune signaling.

However, it is still unclear whether ATM modulates NBS1 dynamics within the micronuclei. First, we examined the number of NBS1-positive micronuclei after inhibiting the ATM kinase activity with Ku55933. We noticed that the number of NBS1-positive micronuclei in response to 6-thio-dG was significantly reduced upon inhibiting ATM kinase activity (**Fig. 5I**). Second, we used the FRAP assay to examine the dynamic binding of NBS1 with micronuclei after inhibiting ATM kinase activity in EGFP-NBS1 transfected cells. We noticed that EGFP-NBS1 binding to chromatin fragments in metaphase was reduced in the presence of an ATM inhibitor (**Fig. 5I**), suggesting that ATM-dependent NBS1 phosphorylation stabilizes the NBS1–chromatin fragment complex.

To verify the influence of ATM kinase activity on NBS1’s diffusion properties, we examined the mobility states of cGAS and NBS1 in the metaphase and telophase stages of BEAS2B cells stably co-expressing TagRFP-cGAS and EGFP-NBS1 cells treated with 20 μM ATM kinase inhibitor for 24 hours by FCS. We noticed that NBS1 mobility increased significantly (p=0.03) in ATM kinase inhibitor-treated metaphase cells (D=27±2.7 μm^2^/s) relative to DMSO-treated metaphase cells (D=19.7±1.1 μm^2^/s; **Fig. 5K**). In contrast, the percentage of fast NBS1 components significantly reduced from ~57% (57.5± 4.1%) in DMSO-treated to ~44% (43.81 ± 5%) in Ku55933-treated metaphase cells (p=0.049; **Fig. 5L**). However, cGAS mobility (D=20±1.1 μm^2^/s and D=20.5±1μm^2^/s in DMSO- and Ku55933-treated, respectively; p=0.1785; **Fig. 5M**) and the percent of fast components (38 ± 2.1% and 39 ± 1.6% in DMSO-and Ku55933-treated, respectively; p=0.47; **Fig. 5N**) in metaphase cells were not significantly changed between DMSO- and Ku55933-treated groups. These observations imply that ATM kinase inhibitor may weaken the steady-state binding of NBS1 to metaphase chromatin fragments. These results imply that the distribution of cGAS molecules changes towards a more stable steady state due to weak chromatin fragment binding characteristics of NBS1 in telophase of ATM kinase inhibitor-treated cells.

### cGAS is not required for NBS1 recruitment to micronuclei, but NBS1 depletion promotes cGAS recruitment to metaphase chromatin and cellular senescence

Although manipulating NBS1 affects the frequency of cGAS-positive micronuclei, it is not clear whether depleting or inhibiting cGAS influences NBS1 binding to micronuclei. Therefore, we evaluated the frequency of NBS1 recruitment to micronuclei in cGAS-dysfunctional cells in response to 6-thio-dG. We found that the number of NBS1-positive micronuclei in shcGAS RNA cells was comparable to that in cells expressing scr-shRNA in response to 6-thio-dG (**Fig. 6A**). Similarly, pharmacological inhibition of cGAS enzymatic activity with RU.521 did not alter the number of NBS1-positive micronuclei in response to 6-thio-dG (**Fig. 6A**). Thus, cGAS is not involved in recruiting NBS1 to micronuclei.

**Figure 6:**
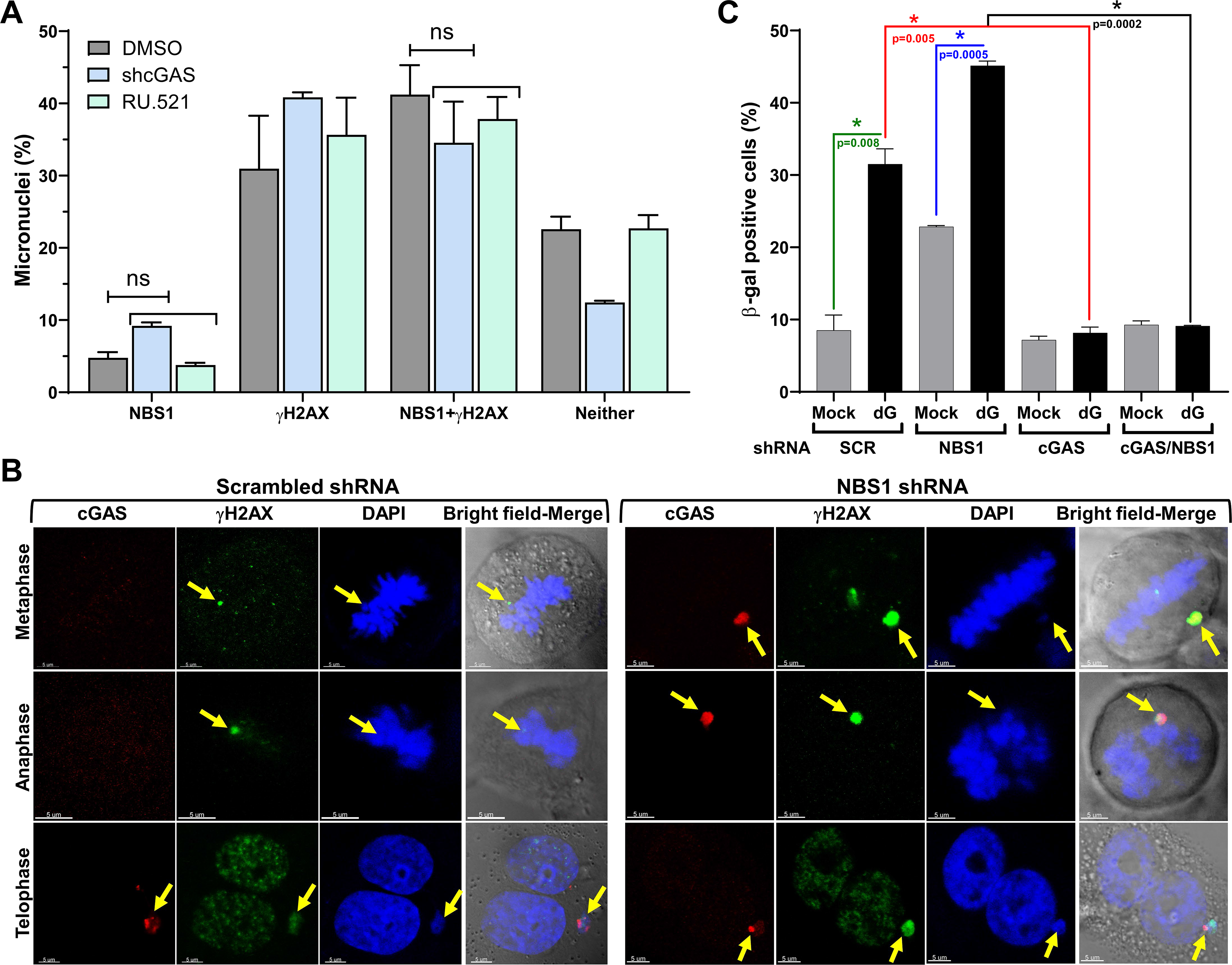
cGAS is not required for NBS1 recruitment to micronuclei but it is needed for cellular senescence in the absence of NBS1. **A.** Depleting cGAS and inhibiting its enzymatic activities do not alter NBS1 binding to micronuclei. Bar graph shows the percentage of micronuclei containing NBS1, γH2AX, both or neither in shcGAS, DMSO- and RU.521-treated BEAS2B cells at 72 hours after 3 μM 6-thio-dG treatment. Bar graph presents the mean and STDEV from 50 different fields of each experimental group. Statistical analysis was performed using student’s t-test. **B.** NBS1-depletion promotes cGAS access to metaphase chromatin fragments in response to 6-thio-dG treatment. Representative images show recruitment of cGAS to chromatin fragments in metaphase, anaphase and telophase stages of BEAS2B-shNBS1 and BEAS2B-SCR cells at 72 hours 3 μM 6-thio-dG treatment. **C.** Co-depleting cGAS and NBS1 abrogates cellular senescence. Bar graph shows percentage of β-gal–positive SCR, NBS1 alone, cGAS alone and NBS1/cGAS shRNAs expressing cells at 10 days after 3 μM 6-thio-dG (dG) treatment. Bar graph presents the mean and STDEV of two independent experiments. Statistical analysis was performed using student’s t-test.

Our FRAP and FCS results clearly indicated that NBS1, but not cGAS, tightly binds to metaphase chromatin fragments and that cGAS starts accumulating onto chromatin fragments during the telophase stage (**Fig. 4**). Hence, it is possible that NBS1 depletion will unblock cGAS’s access to chromatin fragments, leading to cGAS binding to metaphase chromatin, anaphase chromatin, or both. To verify this notion, we examined cGAS recruitment to chromatin fragments in the metaphase and anaphase stages of NBS1-knockdown mitotic phase cells after causing dysfunctional telomeres. Surprisingly, we noticed a clear recruitment of cGAS to chromatin fragments in a major fraction of metaphase and anaphase stages of NBS1-knockdown mitotic phase cells, but not in NBS1-proficient cells (**Fig. 6B**). Collectively, these results reveal that NBS1 is an upstream regulator of cGAS binding to chromatin fragments, and any manipulation of NBS1 impinges on cGAS binding to chromatin fragments, but not vice versa.

cGAS is essential for cellular senescence,^47^ but it is still unclear whether the elevated levels of cellular senescence observed in NBS1-defective cells responding to DNA damage are also mediated by cGAS. To investigate this idea, we co-knocked down NBS1 and cGAS in the same cells by shRNAs, caused genotoxic stress with 6-thio-dG, and counted β-gal–positive cells 10 days later. As shown in **Figure 6C**, the number of β-gal–positive cells in response to 6-thio-dG was significantly lower in co-knocked down cells than in cells depleted of NBS1 alone. Additionally, we noticed that the percentage of β-gal–positive cells in shNBS1-shcGAS cells was similar to that in cells depleted of cGAS alone (**Fig. 6C**). Thus, cGAS is required for causing increased cellular senescence in NBS1-knockdown cells.

### cGAS is not recruited to micronuclei with resected DNA

Next, we set to determine how NBS1 in micronuclei blocks cGAS binding to micronuclei. We speculated that NBS1’s recruitment and stabilization at the DSBs sites of micronuclei alter the DSB ends and that somehow prevent cGAS from binding to all of these micronuclei. Since ΔFHA-NBS1 and ΔATM-NBS1 cells exhibited higher numbers of cGAS-positive micronuclei than WT-NBS1 cells, and because these domains are responsible for recruiting CtIP and RNF20, respectively, to DSBs, we evaluated the activities of CtIP and RNF20 in cGAS binding to micronuclei. First, we verified the recruitment and co-localization of CtIP with cGAS in the micronuclei. As predicted, CtIP was recruited to the micronuclei and clearly co-localized with the γH2AX foci; however, the number of CtIP- positive micronuclei was significantly lower in NBS1-knockdown cells than in NBS1-proficient cells (**Fig. 7A**), which confirms a role for NBS1 in recruiting CtIP to DSB-positive micronuclei.^22^ Interestingly, similar to NBS1, only a minor fraction of cGAS was co-localized with CtIP in micronuclei (**Fig. 7B**). This reduced level of CtIP localization in NBS1-knockdown cells did not result from defective nuclear envelope coating, since CtIP-positive micronuclei were equally positive and negative for Lamin A/C staining (**Fig. 7C**). Second, we examined the recruitment and co-localization of RNF20 with cGAS in the micronuclei. Contrary to our expectation, we rarely detected an RNF20 signal in the micronuclei, either by indirect immunostaining or by live cell imaging (data not shown), which ruled out a possible role for RNF20 in modulating cGAS binding to micronuclei. Thus, CtIP is recruited to micronuclei in an NBS1-dependent manner, and a major fraction of them do not co-localize with cGAS.

**Figure 7:**
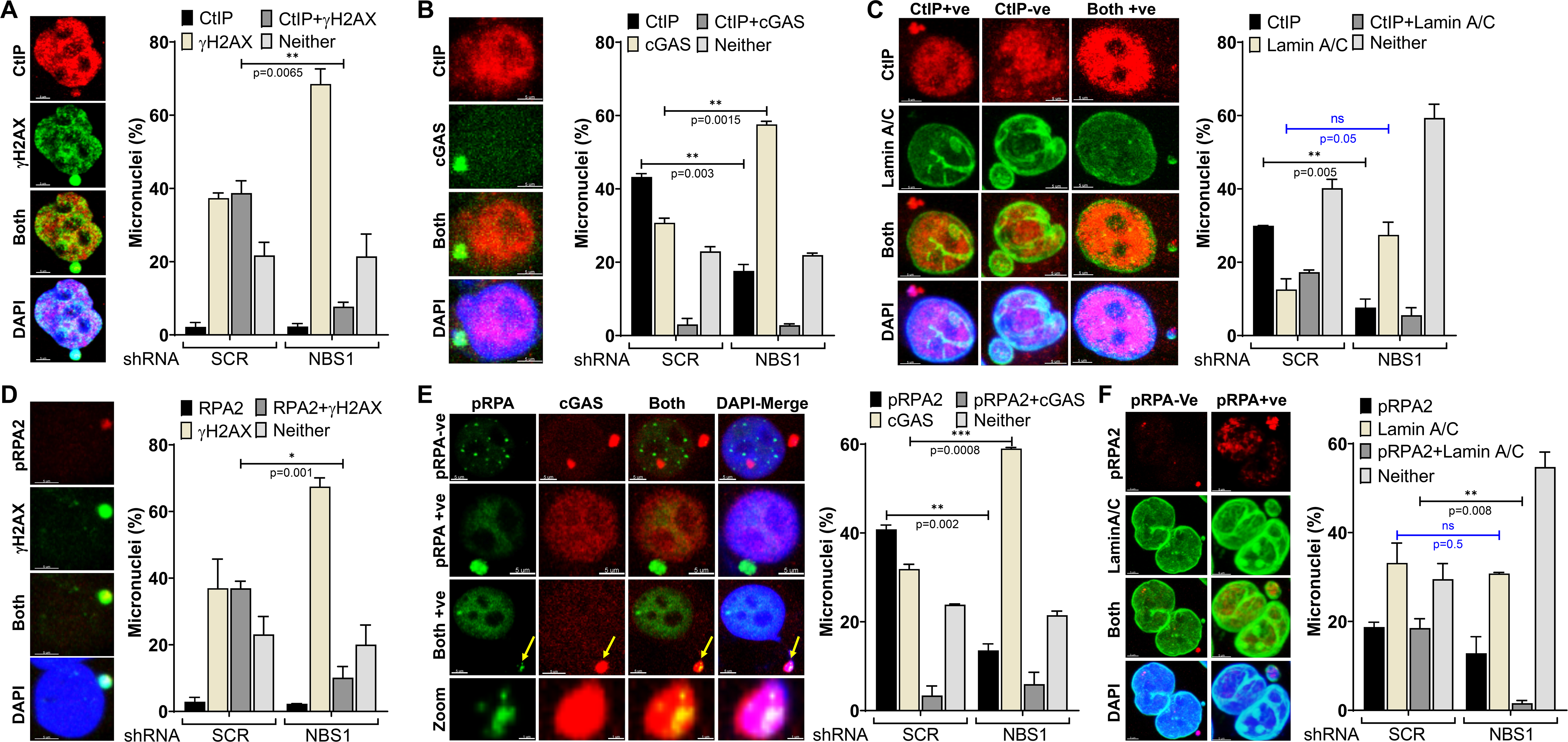
cGAS is not recruited to CtIP- and RPA2-positive micronuclei. **A-C.** CtIP is recruited to micronuclei but does not co-localize with cGAS. Representative images show co-localization of CtIP with γH2AX (**A, left**), cGAS (**B, left**) and Lamin A/C (**C, left**) in the micronuclei. Bar graphs show the frequency of micronuclei harboring CtIP, γH2AX, both or neither (**A, graph**); CtIP, cGAS, both or neither (**B, graph**); CtIP, Lamin A/C, both or neither (**C, graph**) in BEAS2B cells stably expressing either shSCR or shNBS1 RNA at 72 hours after 3 μM 6-thio-dG treatment. Bar graphs present the mean and STDEV from 50 different fields (500-800 cells) from two independent experiments. Statistical analysis was performed using student’s t-test. **D-F.** RPA2 is recruited to micronuclei but does not co-localize with cGAS. Representative images show co-localization of phosphorylated RPA2 (pRPA2) with γH2AX (**D, left**), cGAS (**E, left**) and Lamin A/C (**F, left**) in the micronuclei. Bar graphs show the frequency of micronuclei harboring pRPA2, γH2AX, both or neither (**D, graph**); pRPA2, cGAS, both or neither (**E, graph**); pRPA2, Lamin A/C, both or neither (**F, graph**) in BEAS2B cells stably expressing either shSCR or shNBS1 RNA at 72 hours after 3 μM 6-thio-dG treatment. Bar graphs present the mean and STDEV from 50 different fields (500-800 cells) from two independent experiments. Statistical analysis was performed using student’s t-test.

CtIP resects DSBs in the G1-, S-, G2- and M-phases of the cell cycle, thus generating ssDNA overhangs; this aids RPA2, a surrogate readout for ssDNA, in coating the resected ssDNA.^48–53^ Therefore, we examined whether CtIP recruitment catalyzes DSB resection and leads to the binding of RPA2 onto micronuclei. We detected co-localization of phosphorylated RPA2 (pRPA2) with γH2AX in the micronuclei of shSCR and shNBS1 cells in response to 6-thio-dG (**Fig. 7D**). However, the percentage of pRPA2-positive micronuclei was significantly lower in NBS1-knockdown cells than in shSCR cells (**Fig. 7D**). Like NBS1 and CtIP, we rarely detected co-localization of pRPA2 with cGAS in the micronuclei (**Fig. 7E**). Additionally, we detected pRPA2 signals in both Lamin A/C-positive and negative micronuclei (**Fig. 7F**), which indicates that the lack of co-localization between pRPA2 and cGAS does not result from defective nuclear envelope coating. So, cGAS does not bind to RPA2-positive (i.e. single strand DNA-positive) micronuclei.

### CtIP-catalyzed DNA end resection suppresses cGAS binding to micronuclei

To confirm the relationship between ssDNA ends generated by NBS1-CtIP–initiated DSB resection and cGAS binding to micronuclei, we depleted RNF20 and CtIP, either alone or in combination in BEAS2B cells, and examined cGAS recruitment to micronuclei (**Fig. 8A, inset**). Interestingly, though we detected similar number of micronuclei (**Fig. 8A**), we noticed that more than 80% of micronuclei were positive for cGAS in CtIP-knockdown cells (**Fig. 8B**). However, RNF20 depletion did not significantly alter the number of cGAS-positive micronuclei, as compared with shSCR cells (**Fig. 8B**). As expected, similar to NBS1-knockdown cells, the number of pRPA2-positive micronuclei was significantly lower in CtIP-knockdown cells than in CtIP-proficient cells (**Fig. 8C**). In addition, we observed similar phenotypes in micronuclei generated by ionizing radiation (**Figs. 8D-G**). Ultimately, CtIP-knockdown cells exhibited a significantly increased number of senescent cells after 6-thio-dG and ionizing radiation (**Figs. 8E and G**). Thus, NBS1-CtIP–initiated end-resected micronuclei may not be a preferred binding site for cGAS.

**Figure 8:**
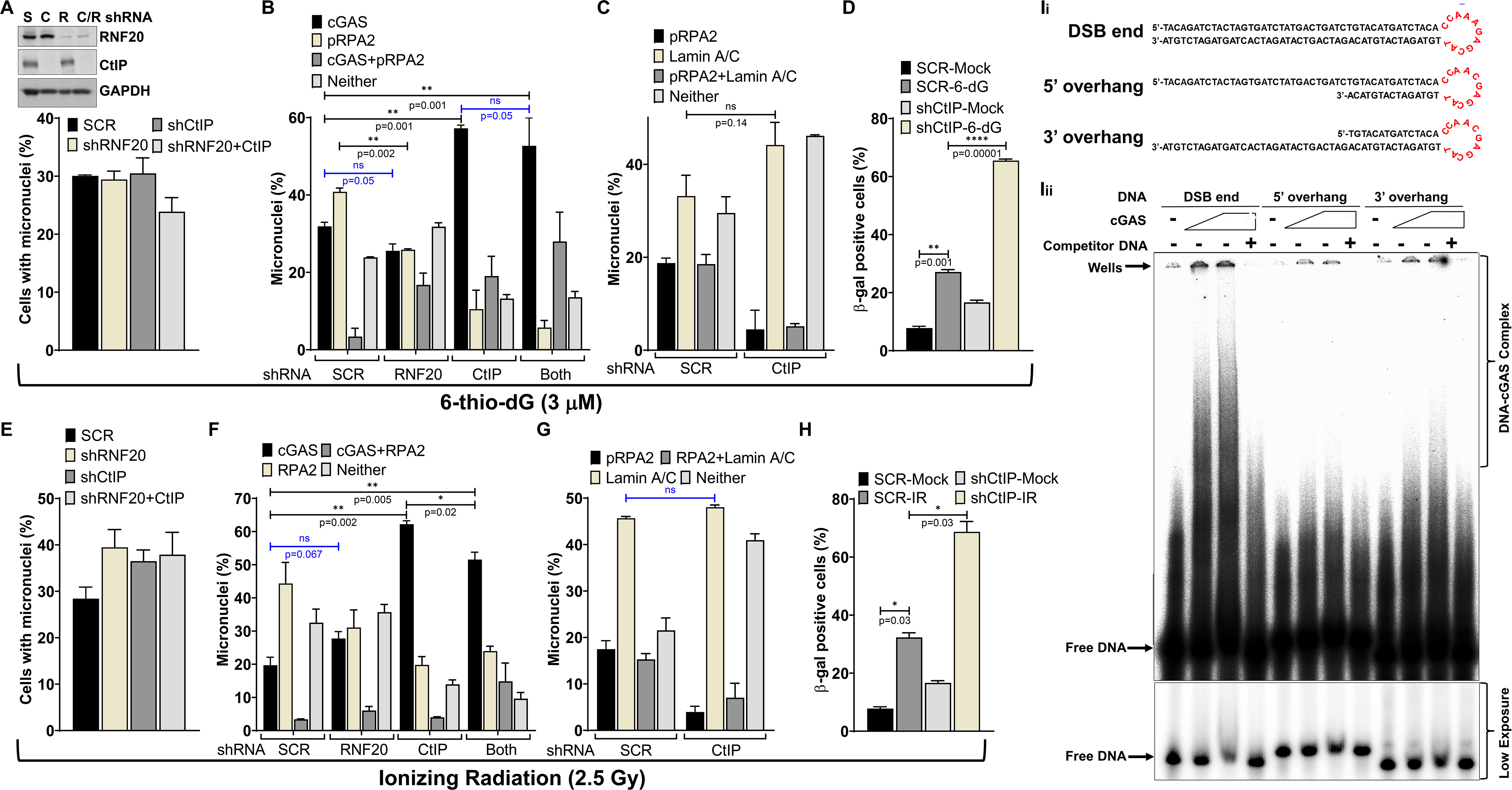
CtIP-mediated DNA end resection attenuates cGAS binding to micronuclei. **A-D.** Downregulation of CtIP enhances cGAS-positive micronuclei and cellular senescence in response to dysfunctional telomeres. Western blots show expression of CtIP and RNF20 in BEAS2B cells stably expressing shSCR (S), shCtIP (C), shRNF20 (R) and shCtIP+shRNF20 (C/R) (**A, inset**). Bar graph shows percentages of micronuclei (**A**); cGAS, pRPA2, cGAS+pRPA2 and neither (**B**); cGAS-, Lamin A/C-, both- or neither-positive (**C**) micronuclei in BEAS2B cells stably expressing scrambled (SCR), shRNF20, shCtIP and shRNF20+shCtIP RNA at 72 hours after 3 μM 6-thio-dG (dG) treatment. Bar graph shows percentage of β-gal–positive cells expressing SCR and CtIP shRNAs at 10 days after 3 μM 6-thio-dG (6-dG) treatment (**D**). Bar graph presents the mean and STDEV of two independent experiments. Statistical analysis was performed using student’s t-test. **E-H.** Depletion of CtIP enhances cGAS-positive micronuclei and cellular senescence in response to DNA lesions induced by ionizing radiation. Bar graph shows percentages of micronuclei (**E**); cGAS, pRPA2, cGAS+pRPA2 and neither (**F**); cGAS-, Lamin A/C-, both- or neither-positive (**G**) micronuclei in BEAS2B cells expressing scrambled (SCR), shRNF20, shCtIP and shRNF20+shCtIP RNA at 72 hours after exposure to 2 .5 Gy. Bar graph shows percentage of β-gal–positive cells expressing SCR and CtIP shRNAs at 10 days after exposure to ionizing radiation (**H**). Bar graph presents the mean and STDEV of two independent experiments. Statistical analysis was performed using student’s t-test. **I.** cGAS binds with double-stranded but not to end-resected DNA substrates. Schematic showing three different DNA structures used for cGAS binding assay (**Ii**). Representative phosphor-image shows cGAS binding with double-stranded but not with end-resected DNA substrates (**Iii**). 5-10 μM cGAS was incubated with 50 femtomole ^32^P labeled DNA substrates in the presence of absence of 100 base pairs cold double stranded DNA (competitor). DNA-protein complex was resolved onto 5% native Poly acrylamide gel electrophoresis and the signal was detected by phosphor imaging.

Yet, it is not clear whether cGAS cannot bind to the end-resected ssDNA. We carried out in vitro experiments by using purified cGAS and DNA substrates that mimic end-resected DNA (**Figs. S3A-C**). Similar to a previous study,^54^ cGAS bound strongly to a 45-base-pairs (bp) interferon stimulatory response element (ISRE) double-strand DNA (dsDNA) substrate (**Figs. 8H and S3E**). However, cGAS binding to the same DNA substrate mimicking either 5’-or 3’-resected DNA was substantially lower (**Fig. 8H**). Collectively, these results provide strong evidence that the coordinated activities of NBS1 and CtIP within micronuclei convert dsDNA ends to ssDNA ends, and this conversion prevents cGAS from binding to all micronuclei.

## Discussion

Previous studies have reported the accumulation of cGAS in nuclear envelope-ruptured micronuclei ^14,29–32^; however, none have provided information on whether cGAS binds to all or to only a sub-set of these micronuclei. Here, we provide both cell biology and biochemical evidence that cGAS is recruited to only a sub-set of micronuclei and that NBS1-CtIP–mediated DSB end resection is responsible for preventing cGAS from binding to all micronuclei. Evidence suggests that cGAS has a higher affinity for nucleosomes than for naked DNA,^55^ and for double-strand DNA (dsDNA) than for single-strand DNA (ssDNA; *K_d_* ~1.5 μM).^6,54^ End resection mediated by NBS1-CtIP and their binding partners not only converts DSB ends into 3’ ssDNA overhangs but also remodels (or relaxes) nucleosomes surrounding the resected DNA.^56^ We detected the recruitment of NBS1, CtIP and RPA2 to micronuclei, which suggests that DNA end resection occurs within these micronuclei.^22^ Additionally, we found that depleting NBS1 and CtIP significantly elevated the number of cGAS-positive micronuclei and that purified cGAS could not bind to a DNA substrate that mimics end-resected DNA substrate. All these observations suggest that the resected ssDNA ends may not be a preferred cGAS binding substrate.

NBS1 was long thought to mainly sense DSBs and transmit DNA damage signaling to regulate genome integrity and cellular homeostasis. Our current findings suggest a novel role for this well-studied DDR factor in limiting cGAS recruitment to micronuclei. NBS1 lacks any known enzymatic activities, but its physical presence on the damaged chromatin and its subsequent recruitment of its binding partners ATM and CtIP to micronuclei choreograph the conversion of dsDNA ends into ssDNA ends, and that conversion blocks cGAS binding to all micronuclei. We provide extensive evidence that: **1.** Both transient and stable knockdown of NBS1 enhances cGAS’s binding efficiency to micronuclei. Because NBS1 recruits CtIP and ATM to DSB sites, NBS1 deficiency causes defects in CtIP-initiated DNA end resection,^22^ which results in the recruitment of cGAS to more micronuclei. **2.** Expression of NBS1 defective in the C-terminal ATM binding domain (delta 705-754 aa) and inhibition of ATM kinase activity increased the number of cGAS-positive micronuclei. ATM kinase activity is important for NBS1 and CtIP phosphorylation, as well as CtIP activation.^22,57^ Defective ATM function attenuates DNA resection and enhances cGAS binding to micronuclei. **3.** Depleting the CtIP-binding FHA-BRCT1/2 domain of NBS1 and CtIP promoted cGAS binding to a major fraction of micronuclei because of the reduced levels of DSB resection within the micronuclei. Overall, these results strongly suggest that the coordinated activities of NBS1 and its binding partners, specifically ATM and CtIP, alter the architecture of micronuclei, which suffices to block cGAS binding to all micronuclei.

CtIP is essential for DSB resection to generate 3’ single-stranded overhangs.^24^ CtIP is recruited to DSB sites by NBS1 through its N-terminal FHA-BRCT1/2 domains.^22^ Furthermore, NBS1-dependent recruitment of ATM phosphorylates CtIP and is also required for its nuclease activity in vitro.^22,58^ Since nucleosome organization forms a barrier for resection, NBS1-CtIP–dependent recruitment of nucleases/helicases (EXO1, DNA2, BLM and WRN) and chromatin remodeling factors (SMARCAD1 and INO80) remodels the chromatin organization and helps to complete the DNA end resection process.^59–63^ Moreover, CtIP-dependent DSB resection is not limited to the S/G2 phases of the cell cycle: DSB resection has also been reported in the G0/G1 and M-phases.^48–53^ Our observations on the recruitment of CtIP to micronuclei and the presence of phosphorylated RPA2 in micronuclei suggest that DSB resection occurs within micronuclei. Additionally, depleting either NBS1 or CtIP not only reduced resection of micronuclei, as indicated by reduced pRPA2-positive micronuclei, but also enhanced cGAS binding to a major fraction of micronuclei. Thus, NBS1-ATM-CtIP–mediated end resection plays a key function in suppressing cGAS interaction with micronuclei. It would be important in the future to understand the roles of DNA end resection nucleases, the chromatin remodelers associated with this process, and the micronuclei harboring difficult to resect DNA ends generated by genotoxic anti-cancer drugs in regulating cGAS activation.

We have combined the data from this study and presented it as a model in **Figure 9**. Following the generation of micronuclei in response to genotoxic stress, NBS1 binds tightly to the damaged micronuclei via NBS1’s FHA domain. Upon binding to γH2AX, NBS1 serves as a docking platform for multiple proteins, including ATM and CtIP. ATM phosphorylates NBS1 and CtIP, in addition to H2AX, which results in both the stabilization of NBS1 and the activation of CtIP. Subsequently, CtIP resects dsDNA ends and converts them into ssDNA ends, which results in their being coated by the ssDNA binding protein RPA2. Because cGAS has a strong affinity for dsDNA but not for ssDNA, these resected DNA may not be a strong binding substrate for cGAS; this prevents cGAS from binding to end resected DSBs of micronuclei. Either after the completion of DSB repair or because of NBS1’s weak binding affinity to certain micronuclei, cGAS binds to a limited number of micronuclei. This regulated binding of cGAS to chromatin fragments initiates innate immune signaling in a limited number of cells, culminating in a premature senescence phenotype. Although the precise mechanism of this process has not yet been verified, our findings will provide a useful approach for further characterizing NBS1 and other DDR factors in regulating cGAS interaction with both cytosolic and nuclear DNA.

**Figure 9:**
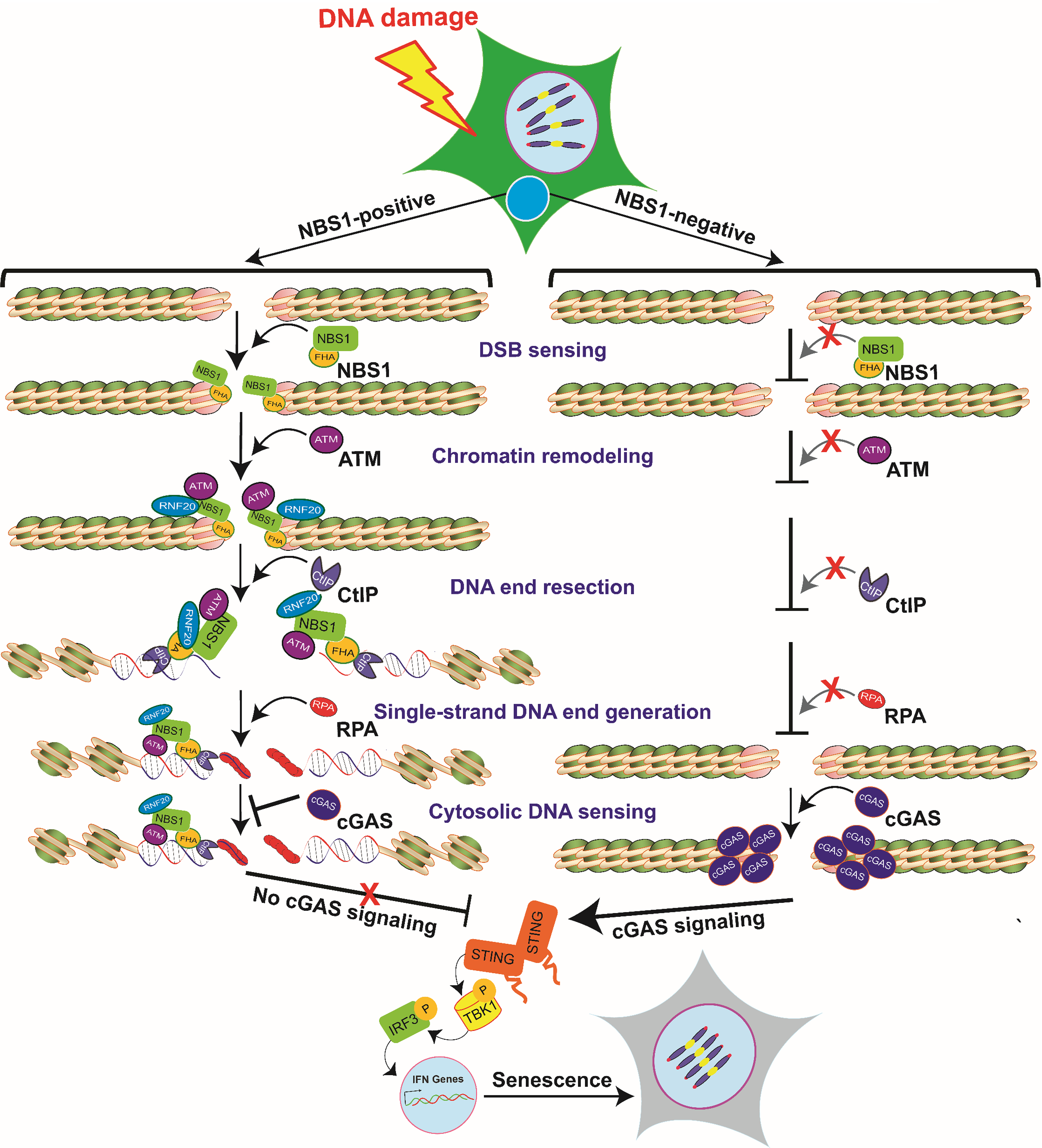
Model depicting the mechanism of NBS1-mediated regulation of cGAS binding to micronuclei and subsequent activation of immune signaling, which culminates in cellular senescence.

In summary, we report a previously unidentified cellular mechanism that regulates cGAS recruitment to micronuclei. We found that cGAS binding is restricted to a limited number of micronuclei in response to genotoxic stress. We have discovered that NBS1’s ability to rapidly and tightly bind to the damaged chromatin and the coordinated activities of its interacting partners ATM and CtIP in processing the micronuclei are the key events responsible for preventing cGAS from binding to all micronuclei. Thus, in addition to its well-studied functions in DNA damage sensing, repair and signaling in response to genotoxic stress, NBS1 acts as an upstream regulator of cGAS recruitment to micronuclei in response to DNA damage.

## Methods

### Cell lines

HT1080, HeLa and BEAS2B cells were obtained from the American Type Culture Collection (ATCC; USA). ATLD and HeLa-shATM cells have been described previously.^37^ NBS (NBS1-deficient) and NBS cells complemented with different NBS1 constructs have been described previously.^37^ HT1080+DD-Cas9/SgTelomere RNA and HT1080-dominant negative (DN)-TRF2 (45 to 453 aa regions of TRF2) cell lines have been described previously.^33^ All cell lines were grown in standard tissue culture conditions at 5% CO_2_ and maintained in Dulbecco modified eagle medium supplemented with 10% fetal bovine serum, 2 mM glutamine and 0.1 mM nonessential amino acids. Mycoplasma contamination was frequently tested by PCR; only Mycoplasma-free cells were used for all our experiments.

To make stable BEAS2B-shcGAS, BEAS2B-shNBS1, BEAS2B-shCtIP, BEAS2B-shRNF20, and BEAS2B-shCtIP+RNF20 RNA cell lines, we infected BEAS2B cells with pooled lentiviral particles carrying either cGAS, NBS1, CtIP, RNF20 or CtIP+RNF20-specific shRNAs (Sigma) and then placed cells under puromycin selection (0.5 μg/ml). To make a stable tetracycline-inducible BEAS2B-shNBS1 RNA cell line, we infected BEAS2B cells with pooled lentiviral particles carrying tetracycline-inducible NBS1-specific shRNAs and then placed cells under G418 selection (500 μg/ml). To stably knock down both cGAS and NBS1 in the same cells, we infected BEAS2B cells with pooled lentiviral particles harboring both shcGAS RNA (Puromycin) and tetracycline-inducible shNBS1 RNA (G418) and selected with puromycin and G418 together. Stable clones were isolated, and the expression of cGAS, NBS1, CtIP and RNF20 was verified by western blotting.

To make a stable doxycycline-inducible TagRFP-Flag-cGAS and mAmetrine Lamin B10_BEAS2B cell line, we infected BEAS2B cells with lentivirus carrying tetracycline-inducible TagRFP-Flag-cGAS-Hyg^R^, then placed them under hygromycin selection (150 μg/ml). We isolated stable clones and treated them with Doxycycline (0.25-1 µg/ml) for 72 hours, then verified the expression and micronuclei localization of cGAS by both western blotting using anti-FLAG antibodies and live cell imaging microscope. Subsequently, we used Lipofectamine 2000 to transfect BEAS2B+TagRFP-Flag-cGAS cells with mAmetrine-Lamin B10 _G418^R^, then placed them under G418 selection (500 μg/ml). Finally, we verified the localization of mAmetrine-Lamin B10 by using a live cell imaging microscope.

### DNA manipulation and construction of the expression vectors

Standard molecular biology procedures were used to make all mammalian expression plasmids. A panel of human cGAS-specific shRNAs (TRCN0000128706, TRCN0000128310, TRCN0000149984, TRCN0000146282, and TRCN0000150010), human NBS1-specific shRNAs (TRCN0000295898, TRCN0000040137, TRCN0000040133, TRCN0000012671 and TRCN0000012672), human CtIP-specific shRNA (Sigma, TRCN0000318738, TRCN0000005403, and TRCN0000005405) and human RNF20-specific shRNA (Sigma, TRCN0000033876, TRCN0000033877, and TRCN0000033878) was purchased from Sigma. To make tetracycline-inducible shNBS1 RNA constructs, we annealed shNBS1-F1/F2/F3 and shNBS1-R1/R2/R3 primers (**Table S1**) in annealing buffer (100 mM NaCl, 10 mM Tris-HCl, pH=7.4), then ligated them into AgeI-EcoRI sites of a Tet-pLKO-neo vector.

To make doxycycline-inducible TagRFP-Flag-cGAS constructs, we PCR amplified TagRFP-Flag-cGAS (Addgene) by using RFP-cGAS-XbaI-F and cGAS-XhoI-R primers (**Table S1**). PCR-amplified fragments were digested with XbaI and XhoI enzymes (NEB) and cloned into the NheI and SalI sites of pCW57-MCS1-P2A-MCS2 (Hygro) vector (Addgene).

Full-length NBS1 (FL-NBS1) has been described previously.^37^ To make the NBS1_1-200 aa_ region of NBS1, we PCR amplified the 1-200 aa region by using FL-NBS1 plasmid and NBS1-1F and NBS1-200R primers (**Table S1**). PCR-amplified fragments were digested with NheI and SalI enzymes (NEB) and cloned into the NheI and SalI sites of a pCW57.1 vector. We confirmed the sequence of all constructs prior to use. All FLAG-tagged protein constructs are N-terminal fusions.

### Addgene plasmids used in this study

mAmetrine-LaminB1-10 was a gift from Michael Davidson (Addgene plasmid # 56547); pTRIP-CMV-tagRFP-FLAG-cGAS was a gift from Nicolas Manel (Addgene plasmid # 86676); pCW57.1 was a gift from David Root (Addgene plasmid # 41393); pCW57-MCS1-P2A-MCS2 (Hygro) was a gift from Adam Karpf (Addgene plasmid # 80922); Tet-pLKO-puro-Scrambled was a gift from Charles Rudin (Addgene plasmid # 47541); Tet-pLKO-neo was a gift from Dmitri Wiederschain (Addgene plasmid # 21916); pCW-Cas9 was a gift from Eric Lander & David Sabatini (Addgene plasmid # 50661).

### Lentiviral production

293T cells were co-transfected with highly purified expression plasmids and pPLP2/pLP1/pVSVG by using either lipofectamine 2000 (Invitrogen), according to manufacturer’s instructions, or the calcium phosphate method described by Kwon et al.^64^ We collected cell culture supernatant containing viral particles 72 hours after the transfection, filtered it through 0.22 microns PVDF membrane filters (Millipore), and used the flowthrough to infect the cells.

### Chemicals

We used 6-thio-dG (Sigma, #1296), ATM inhibitor (Sigma, #925701-46-8), doxycycline (Sigma, #D9891), cGAS inhibitor RU.521 (AOBIOUS, #37877), Shield1 (Cheminpharma, #S1-0005), Puromycin (AdipoGen), Hygromycin (Invitrogen), G418 (Sigma Aldrich), gemcitabine (Selleck Chemical, #S1149), hydroxyurea (Sigma Aldrich, # H8627) and aphidicolin (EMD Millipore, #178273).

### Doxycycline and Shield1 treatment

**a.** To induce DSBs in the telomeric DNA of HT1080-sgTelomere-DD-Cas9 cells, we co-treated cells with 500 ng/ml doxycycline (Sigma) and 1 μM Shield1 (Cheminpharma) for 24 hours. Cells were then washed three times with warm PBS and allowed to recover in regular growth medium; samples were collected after 24-72 hours. **b.** To induce DN-TRF2 expression, we treated HT1080-DN-TRF2 cells with 1000 ng/ml doxycycline (Sigma) for 72 hours. Cells were then washed three times with warm PBS and allowed to recover in regular growth medium; samples were collected after 24-74 hours post-DOX withdrawal (w/d) time points. **c.** To deplete NBS1, we treated cells expressing BEAS2B-shNBS1 RNA with 500 ng/ml doxycycline (Sigma) for 72 hours. Cells were then treated with 6-thio-dG (3-5 μM) in the presence of 500 ng/ml doxycycline (Sigma), washed three times with warm PBS and allowed to recover in regular growth medium containing 500 ng/ml doxycycline; samples were collected at different post-DOX withdrawal (w/d) time points. **d.** To induce TagRFP-cGAS, we treated BEAS2B_TagRFP-Flag-cGAS_mAmetrine Lamin B10 cells with 250 ng/ml doxycycline (Sigma) for 72 hours. Samples were collected by mitotic shake-off, allowed to attach on glass-bottomed petri dishes (MatTeK) and used for live cell imaging.

### 6-thio-dG treatment

Twenty-four hours after plating, we treated cells with 2-5 μM 6-thio-dG for 72 hours, washed three times in warm PBS, and then cultured in drug-free medium for an additional 0-15 days. Samples were collected at different post–6-thio-dG recovery times.

### cGAS inhibitor (RU.521) treatment

Cells were treated with 1 μM RU.521 dissolved in DMSO for the entire duration of the experiment. Fresh drug was added to the medium every 48 hours for continuous inhibition of cGAS.

### ATM inhibitor (Ku55933) treatment

Cells were treated with 10-20 μM Ku55933 dissolved in DMSO for the entire duration of the experiment. Fresh drug was added to the medium every 24 hours for continuous inhibition of ATM kinase activity.

### Ionizing radiation

We exposed exponentially growing cells to 2-5 Gy γ-rays by using a γ-irradiator (Mark 1 irradiator, JL Shepherd & Associates), as described previously.^65^

### Cell extracts and western blotting

We prepared whole-cell extracts by suspending cell pellets in RIPA buffer containing protease (PMSF, aprotinin, leupeptin, pepstatin A, dithiothreitol; all at 1:1000 dilutions) and phosphatase (Na_3_VO_4_ at 1:500 and NAF at 1:200 dilution) inhibitors on ice for 30 minutes, followed by centrifugation at 14,000 RPM for 30 minutes at 4°C to remove insoluble material. Whole-cell extracts (30 to 100 μg) were resolved by 6-12% SDS-PAGE, transferred onto Polyvinylidene difluoride membranes and incubated with antibodies of interest.

### Antibodies used in this study

All the primary antibodies used in this study are detailed in **Table S2**, including vendors’ information, application and the dilution conditions. Manufacturers’ validation criteria were used for the application of all antibodies. For western blotting, HRP-conjugated goat anti-rabbit and anti-mouse secondary antibodies were purchased from BioRad and used at 1:1000 dilution in 5% BSA or milk. For indirect immunofluorescence staining, fluorescent-conjugated secondary antibodies Alexa-488, Alexa-555 and Alexa-633 were purchased from Molecular Probes (Invitrogen) and used at 1:1000 dilution.

### Indirect immunostaining

Approximately 0.5-1 × 10^5^ cells were plated in a six-well plate containing cover glasses and incubated for 36 hours. Cells were treated with various chemicals for different time periods, as described above. Cells were fixed with 4% PFA for 20 minutes at room temperature at different post-treatment times and subjected to indirect immunofluorescence, as described previously. Briefly, cells were permeabilized in Triton X-100 (0.5% in PBS) on ice for 5 minutes, washed three times with PBS, incubated in blocking solution (5% goat serum in PBS) at room temperature for 60 minutes, and then incubated with primary antibodies (diluted in 5% goat serum) at room temperature for another 3 hours or at 4°C overnight. Then, cells were washed with wash buffer (1% BSA in PBS), incubated with appropriate secondary antibodies (1:1000 in 2.5% goat serum, 1% BSA, and PBS) at room temperature for 60 minutes, washed five times with 1% BSA, and mounted with mounting medium containing DAPI (Vectashield).

### Image acquisition and foci dissolution kinetics assay

Images were captured using an LSM 510 Meta laser scanning confocal microscope with a 63X1.4 NA Plan-Apochromatic oil immersion objective. Images were taken of z-sections (15-20 sections) at 0.35-μm intervals using the 488-nm (EGFP and Alexa 488), 543-nm (Alexa 555), 633-nm (Alexa 633) and 405-nm (for DAPI) lasers. The tube current of the 488-nm argon laser was set at 6.1 A. The laser power was typically set to 3-5% transmission with the pinhole opened to 1-2 Airy units. To enumerate individual and co-localization of proteins in the micronuclei, we assembled the z-sections by using the Imaris software and analyzed them as described previously.^33^ We quantified individual and co-localized proteins from images of 50-1000 cells per time point from two to three independent experiments.

### Micronuclei imaging and quantification

Cells fixed with 4% PFA were mounted with mounting solution containing DAPI (Vectashield). Images were acquired using an Axio-Invert (Zeiss) microscope using the DAPI channel, and the exposure time ranged from 400 to 500 μ seconds per frame. We used ImageJ (NIH) to calculate micronuclei in a blinded fashion in 100-1000 cells per experimental condition in two to three independent experiments.

### Senescence assay

We subjected freshly fixed samples to β-galactosidase (β-gal) staining by using either β-gal staining kit (Cell Signaling), following manufacturer’s instructions, or homemade β-gal staining solution, as described previously.^33^ We acquired random images by using the 10X objective of a KEYONCE Microscope in a blinded manner. We counted in a blinded fashion the number of β-gal–positive cells in 20-30 random fields consisting of 1000-5000 total cells per experimental condition in two to three independent experiments.

### Quantitative real-time polymerase chain reaction (qRT-PCR)

We synthesized cDNA from 1-3 μg of total RNA by using SuperScript III Reverse Transcriptase (18-080-051; Fischer Scientific) in a total volume of 20 µl, according to manufacturer’s instructions. We subjected the cDNA to qRT-PCR for several genes by using the primer sets (**Table S3**), CFX96 Touch Real-Time PCR Detection System (Bio Rad) and iTaq Universal SYBR Green Supermix (Bio Rad; #1725121), according to manufacturer’s instructions. Relative gene expression was determined by the ΔΔCT method. The difference in cycle times, ΔCT, was determined as the difference between the tested gene of interest and the reference housekeeping β-actin gene. We then obtained ΔΔCT by finding the difference between the groups. The fold change (FC) was calculated as FC=2^−ΔΔCT^. All primers were purchased from Invitrogen. qRT-PCR assays were carried out in triplicate for each sample, and the mean value was used to calculate mRNA expression levels.

### Live cell imaging

Live cell imaging was carried out as described previously.^66^ Briefly, BEAS2B cells stably expressing mAmetrine-LaminB10 (G418^R^) and Tag-RFP-cGAS (Hygromycin^R^) were transfected with EGFP-NBS1 and then treated with 250 ng/ml DOX for 72 hours. The mitotic cells were collected by mitotic shake-off and then seeded onto round 25 mm cover glasses (Electron Microscopy Sciences) and placed onto the cell-holding chamber of an LSM510-Meta confocal microscope equipped with a CO_2_ incubator and a heating chamber. To identify micronuclei and cells in different stages of the mitotic phase, we incubated cells with Hoechst (Invitrogen) in the cell holding chamber for 5-10 minutes, washed the cells with regular growth medium and then subjected them to live cell imaging. We used 405- (Hoechst), 437- (mAmetrine-Lamin B0), 488- (EGFP-NBS1) and 514- (Tag-RFP-cGAS) nm lasers to capture the images in four different channels. Cells in 6-8 fields were imaged every 10-15 minutes and continuously followed for 12-16 hours.

### Fluorescence Recovery After Photobleaching (FRAP)

FRAP was carried out as described previously.^66^ Briefly, we used cells and experimental conditions similar to those described in the live cell imaging section. The NBS1 and cGAS fluorescence signal in half (or less than half) of the micronuclei was photobleached with green (EGFP) and red (Tag-RFP) lasers, respectively. The FRAP signal was calculated as described previously.^66^

### Fluorescence Correlation Spectroscopy (FCS)

FCS was carried out as described previously.^40^ Briefly, we used cells and experimental conditions similar to those described in the live cell imaging section. FCS was performed with a Zeiss ConfoCor3 system and a C-Apochromat 40X numerical aperture 1.2 water immersion objective, as previously described.^40^ EGFP-NBS1 and Tag-RFP-cGAS were excited with 408 and 514 nm lasers, respectively. The normalized ACF (G (τ)) of fluorescence fluctuations was calculated using Zeiss ConfoCor2 software as described previously ^40^.

### ELISA

We collected cell culture supernatant at different post-treatment times and centrifuged at 800 g for 5 minutes at room temperature to remove any cell debris. We then measured cytokine levels by ELISA using the Biolegend human IL-6 (Cat# 430504) kit, as per the manufacturer’s instructions and as described previously.^67^

### Expression and purification of human cGAS

Human cGAS-expressing plasmid construct pET28a-His-hcGAS was a kind gift from Dr. Li.^54^ Purified plasmid was transformed into BL21(DE3)-competent cells and selected using Kanamycin (50 μg/ml). A single bacterial colony was grown in regular LB medium containing 50 μg/ml Kanamycin at 16°C to an optical density of 0.25 at 580 nM. Protein expression was initiated by IPTG (500 ng/ml) at 16°C overnight. cGAS purification was carried out as described previously.^68^ Briefly, BL21(DE3) cell pellets were resuspended and lysed by sonication in 25 mM Hepes-NaOH pH 7.9, 5% glycerol, 20 mM Imidazole, 300 mM NaCl, and 0.2 mM PMSF. Lysate was clarified by centrifugation at 100,000 x g for 1 hr. Clear lysate was loaded into a 5 mL HisTrap HP (GE Healthcare) column, and, after the column was washed with 25 mM Hepes-NaOH pH 7.9, 5% glycerol, 25 mM Imidazole, 300 mM NaCl, and 0.2 mM PMSF, cGAS was eluted with a gradient of 25-500 mM Imidazole. Fractions containing cGAS were combined and loaded into 1 mL Heparin (GE Healthcare). After being washed with 25 mM Hepes-NaOH pH 7.9, 5% glycerol, 0.1 mM EDTA, 500 mM NaCl, 1 mM DTT, and 0.2 mM PMSF, cGAS was eluted with a gradient of 0.5-1 M NaCl. Fractions containing cGAS were combined and loaded into Superdex 200 Increase 10/300 GL (GE Healthcare) in 25 mM Hepes-NaOH pH 7.9, 5% glycerol, 0.1 mM EDTA, 150 mM NaCl, 1 mM DTT, and 0.2 mM PMSF.

### Gel shift analysis

This was performed as described previously.^69^ Briefly, samples were assembled in 20 μl reactions containing 10 mM Hepes-NaOH pH 7.5, 100 mM NaCl, 1 mM DTT, 1 mM MgCl_2_, 20 μg/mL BSA, 25-50 femtomole DNA, and cGAS. After incubating at 25°C for 30 minutes, we stopped the reactions by adding 2 μl of 50% glycerol, 0.05% bromophenol blue, 0.05% xylene cyanol, and 20 mM EDTA. Samples were loaded onto 6% native polyacrylamide gel (19:1) in 0.5x TBE. Gel electrophoresis was performed at room temperature.

### Statistics

Statistical analysis of data was performed using GraphPad Prism Software (version 8.4.2). Two tailed unpaired Student’s t tests and two-way analysis of variance (ANOVA) were used for statistical analyses, and unless otherwise noted, all results are representative of at least two independent biological experiments and are reported as the mean ± standard deviation. GraphPad Prism (version 8.4.2) was used to create the graphs.

### Data availability

The data that support the findings of this study are available from the corresponding author upon reasonable request.

## Acknowledgements

This work was supported by the National Institutes of Health grants R01AG053341 (to A.A.) and R01HL115275 (to H.A.S) and the Cancer Prevention and Research Institute of Texas grant RP190435 (to A.A. and H.A.S). We thank Dr. Jonathan Feinberg for editing this manuscript.

## Authors’ contributions

S.A., S.M., S.B., and A.A. designed the experiments. S.A, S.M., S.B., S.K., J.R.O. and A.A. performed experiments and analyzed data. D.S. performed imaging and quantified micronuclei. S.K. carried out western blotting; S.A. performed all microscopic imaging; S.M. generated cell lines, performed qRT-PCR and immunostaining and analyzed the results; J.R.O. purified cGAS and performed gel shift analysis. H.A.S. and G.L. provided reagents used in the studies, reviewed the manuscript and provided feedback/discussion. S.M., S.B. and A.A. directed the studies, and A.A. wrote the manuscript.

